# Rapid kinetics of lipid second messengers controlled by a cGMP signalling network coordinates apical complex functions in *Toxoplasma* tachyzoites

**DOI:** 10.1101/2020.06.19.160341

**Authors:** Nicholas J. Katris, Yoshiki Yamaryo-Botte, Jan Janouškovec, Serena Shunmugam, Christophe-Sebastien Arnold, Annie S. P. Yang, Alexandros Vardakis, Rebecca J. Stewart, Robert Sauerwein, Geoffrey I. McFadden, Christopher J. Tonkin, Marie-France Cesbron-Delauw, Ross. F. Waller, Cyrille Y. Botte

**Affiliations:** ApicoLipid Team, Institute for Advanced Biosciences, CNRS UMR5309, Universite’ Grenoble Alpes, INSERM U1209, Grenoble, France; Department of Pharmacy, University of Oslo Norway; Radboud University Medical Center, PO Box 9101, 6500 HB Nijmegen, The Netherlands; Department of Biochemistry, University of Cambridge, Cambridge, UK; The Walter and Eliza Hall Institute of Medical Research, Parkville, Melbourne, Victoria, Australia; McFadden Laboratory, School of Biosciences, University of Melbourne, Melbourne, VIC 3010, Australia; Department of Medical Biology, The University of Melbourne, Melbourne, Victoria, Australia

## Abstract

Host cell invasion and subsequent egress by *Toxoplasma* parasites is regulated by a network of cGMP, cAMP, and calcium signalling proteins. Such eukaryotic signalling networks typically involve lipid second messengers including phosphatidylinositol phosphates (PIPs), diacylglycerol (DAG) and phosphatidic acid (PA). However, the lipid signalling network in *Toxoplasma* is poorly defined. Here we present lipidomic analysis of a mutant of central flippase/guanylate cyclase TgGC in *Toxoplasma*, which we show has disrupted turnover of signalling lipids impacting phospholipid metabolism and membrane stability. The turnover of signalling lipids is extremely rapid in extracellular parasites and we track changes in PA and DAG to within 5 seconds, which are variably defective upon disruption of TgGC and other signalling proteins. We then identify the position of each protein in the signal chain relative to the central cGMP signalling protein TgGC and map the lipid signal network coordinating conoid extrusion and microneme secretion for egress and invasion.

## Introduction

The phylum Apicomplexa includes infectious agents of major human and animal diseases such as malaria and toxoplasmosis, which create a massive global burden for human health (World Health Organization, 2018). These parasitic protists are obligate intracellular parasites of human cells, which means that they must invade a human cell in order to divide, persist and cause disease. Central to this process of host cell invasion is a unique structure at the apex of the parasite cell termed the apical complex, which also defines the name of the Apicomplexa phylum. The core of the apical complex is made up of tubulin based cytoskeletal components forming a circular apical polar ring, to which are connected a series of sub-pellicular microtubules which cascade down the length of the parasite body. Apicomplexan parasites also bear an arsenal of secretion factors including microneme and rhoptry proteins, which are secreted through the apical complex to allow invasion and subsequent egress. Interestingly, the apical complex bears an additional tight-knit cytoskeletal structure, the conoid, best studied in *Toxoplasma*. This conoid is connected to the apical polar ring and can extrude and retract from as part of the apical complex (Graindorge et al. 2016). However, the conoid is less conspicuous, or even proposed to be absent from *Plasmodium* species (Wall et al. 2016). The apical complex (bearing or not the conoid) is also present in some photosynthetic relatives of Apicomplexa, such as *Chromera velia* (Oborník et al. 2012; 2011). The conoid is presumed to have a function in parasite invasion as it is typically extruded in extracellular stages, and retracted during intracellular replication (Graindorge et al. 2016; Katris et al. 2014), although its precise function is not fully resolved.

Parasite invasion of a host cell requires the release of secretion factors such as micronemes and rhoptries, both of which are well characterized. Micronemes are essential for attachment to the host surface and required for active motility in both host cell invasion and egress (Bullen et al. 2016). Similarly, rhoptries are required to form the junction between the parasite and host cell during the host entry, an invasion-specific event which is dispensable for egress (Mueller et al. 2013). The signalling mechanism behind microneme secretion is well documented and is regulated by the highly conserved eukaryotic Protein Kinase G (PKG) in both *Plasmodium* zoite stages and *Toxoplasma* tachyzoites (Brochet et al. 2014; Wiersma et al. 2004). PKG has been shown to be essential both in every stage of the *Plasmodium* life cycle including blood, mosquito and liver stages, and in *Toxoplasma* asexual tachyzoites, and is critical for microneme secretion to facilitate invasion and egress in both parasite species (Brochet et al. 2014). PKG is a highly conserved signalling protein, which is typically present in flagellated eukaryotes, where it organizes flagellar movements (e.g., in ciliated mammalian cells and the free-swimming alga *Chlamydomonas reinhartii*), and typically absent in non-flagellated eukaryotes such as higher plants and fungi (Johnson and Leroux 2010). Therefore, the finding that PKG regulates apical complex secretion is consistent with the association of the apical complex with proteins homologous to algal flagellar root components (Francia et al. 2012).

In metazoans such as human cells and *C. elegans*, calcium signalling is regulated by the well characterized G-Protein Coupled Receptors (GPCRs), GPCR activation triggers cAMP production and calcium release which activates Protein Kinase C (PKC) (Xu and Jin 2015). In particular, GPCRs trigger the highly conserved signalling cascade to breakdown lipid second messengers phosphatidylinositol-biphopshate (PIP2) into diacylglycerol (DAG) and inositol triphosphate (IP3), a fundamental eukaryotic signalling process, wherein the cytosolic IP3 activates ER-resident calcium channels to initiate a stress response (Bahar, Kim, and Yoon 2016). However, canonical metazoan GPCRs are absent in Apicomplexa which instead rely on activation of guanylate cyclase and cGMP signalling to activate intracellular calcium release (Stewart et al. 2017; Brochet et al. 2014) for invasion and egress processes. Furthermore, apicomplexan parasites possess a set of plant-like CDPKs which regulate secretion and motility in response to IP3-dependent calcium release for life-stage specific functions (Brochet et al. 2014; McCoy et al. 2012), some of which have roles at intersections with the cGMP pathway (Brochet et al. 2014; McCoy et al. 2012). Other core eukaryotic signalling elements in the cGMP pathway are absent in Apicomplexa including calcium-gated ion channels and soluble and membrane forms of guanylate cyclase. Instead, apicomplexans possess an unusual alveolate-specific dual-domain phospholipid transporter/cGMP cyclase protein (TgGC) fusion protein (Johnson and Leroux 2010; Yang et al. 2019). Recently in *Toxoplasma*, TgGC has been shown to be critical for activation of TgPKG to trigger microneme secretion (Brown and Sibley 2018; Yang et al. 2019; Bisio et al. 2019). Similarly, *Plasmodium* species possess two GCs, type a and type b (GCa + GCb), and the *Plasmodium yoelli* protein PyGCb has been shown to have an apical localization in sporozoites, critical for ookinete traversal of the mosquito midgut (Gao et al. 2018). PyGCa however has a role in blood stage gametocytogenesis and its roles in intraerythrocytic blood stage and mosquito stage have yet to be determined (Jiang et al. 2020).

The function of the P-Type ATPase domain of TgGC is poorly understood but is predicted to have a role in lipid trafficking across membranes, or possibly of ion transport such as K^+^ or Ca^2+^ (Yang et al. 2019; Bisio et al. 2019). However, several studies have concluded that both domains are essential for TgGC function and that the ATPase domain likely links a metabolic function to a signalling response (Brown and Sibley 2018; Yang et al. 2019), although extensive lipidomic analysis in TgGC-depleted cells is lacking. Exactly what the substrate of the ATPase domain might be is uncertain although it has been shown that cells with disrupted TgGC are less responsive to changes in pH while disruption of cofactors of TgGC, including TgCDC50, leads to parasites that can still egress in response to changes in pH (Bisio et al. 2019), suggesting that TgGC is the primary responder to external pH signals. This change in pH is thought to be mediated at least in part by phosphatidic acid (Bisio et al. 2019), which due to its sole phosphate headgroup can bear variable charge through variable binding of hydrogen groups from the external environment (Abramson et al. 1964; van Paridon et al. 1986).

Despite evidence of TgGC being involved in PA signalling, there is no evidence yet for any phosphatidylinositol (PI) during signalling in *Toxoplasma* invasion or egress. This is of particular importance as PI has also been shown to act as a pH biosensor in the Golgi of fibroblasts (Shin et al. 2020). Moreover, despite the existence of a putative PI-PLC in *Toxoplasma* (Bullen et al. 2016), diacylglycerol is still a poorly understood second messenger in *Toxoplasma*. Here we extend on the cGMP signalling model to map out the lipid signalling network. We perform exhaustive lipidomic analysis of TgGC-depleted parasites and establish TgGC as the central regulator of lipid signalling controlling rapid egress and invasion of tachyzoites. Loss of TgGC impacts conoid extrusion, the regulation of which is tightly integrated into the cGMP signalling network controlling apical secretion of micronemes. We further show that TgCDPK1, TgPKG, and TgRNG2 are some of the key proteins involved in this cGMP-dependent lipid signalling network. Crucially, we identify putative homologues of proteins involved in the phosphatidylinositol cycle including a set of putative phospholipid transporters with putative phosphatidic acid binding (PH domain) and oxysterol binding proteins domains (OSBPs), to maintain phospholipid homeostasis (Kim et al. 2015; Shin et al. 2020) and that this is likely regulated by the cGMP signalling network controlled by TgGC.

## Results

### TgGC has a dynamic localization

TgGC is the only flippase, (lipid transporters that flip lipids across membranes) and an unusual Alveolate-specific dual domain flippase-guanylate cyclase fusion protein (**Figure S1**). Previous studies all showed that TgGC is primarily found at the apical cap (Yang et al. 2019; Brown and Sibley 2018; Bisio et al. 2019), but it is also present in other secondary locations including an ER-like compartment (Brown and Sibley 2018), and the residual body (Bisio et al. 2019). This mixed distribution is consistent with a recent hyperLOPIT (Localization of Organellar Proteins by Isotopic Tagging) study that did not assign TgGC to any one location although did indicate a dynamic distribution including the Golgi (Barylyuk et al., n.d.) (www.toxodb.org). After imaging HA-tagged parasites, we found TgGC was localized primarily to the apical cap but we also saw TgGC at what appears to be a Golgi/ER-like compartment just anterior to the nucleus (**Figure 1A**), consistent with the hyperLOPIT signal. Our data supports that TgGC has a very dynamic localization in tachyzoites, with the apical cap as the primary localization, but also with prominent secondary localizations at Golgi/ER-like compartments.

**Figure 1:**
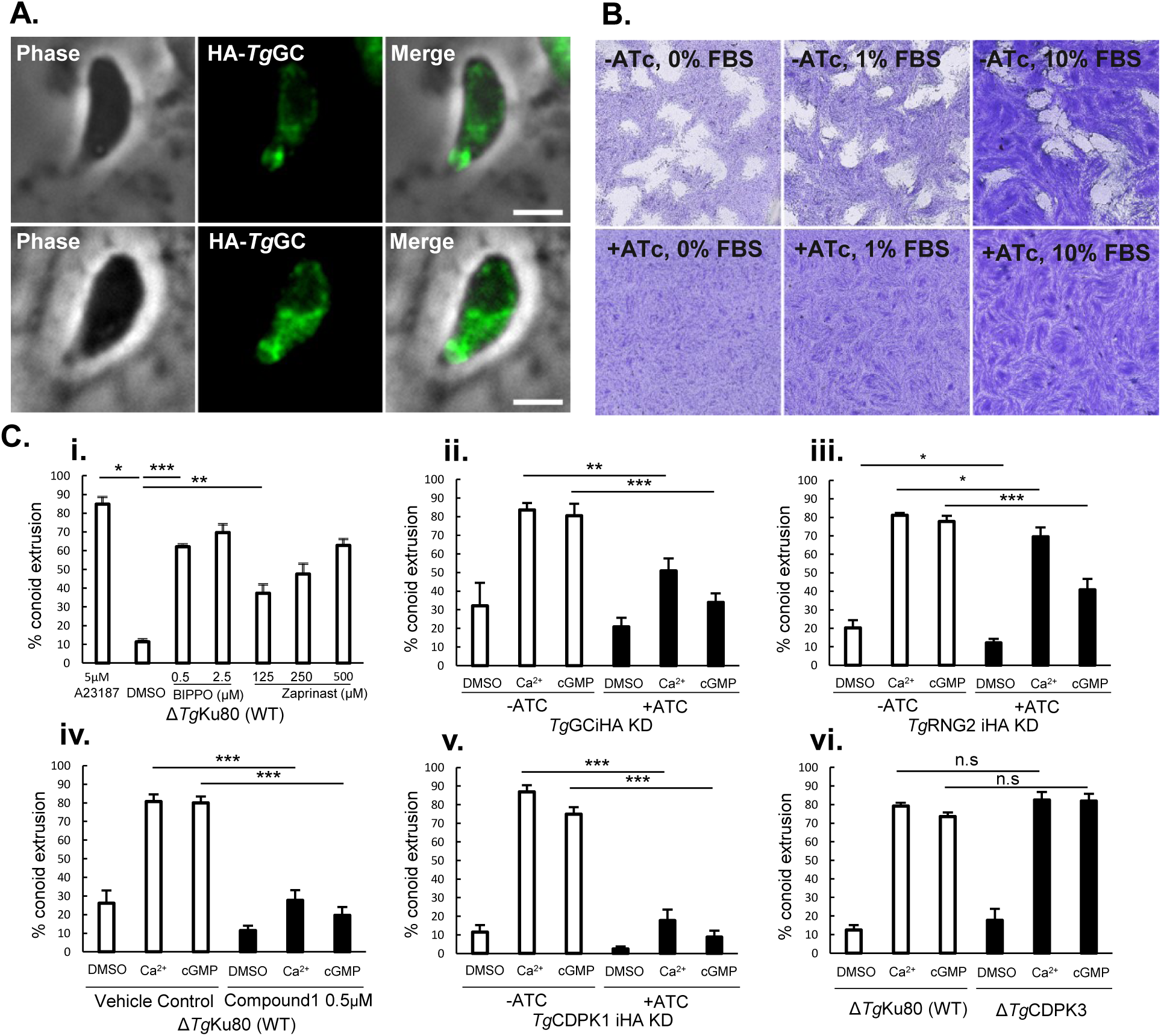
cGMP signalling regulates conoid extrusion. **A** Localization of TgGC (TGGT1_254370) to the apical cap and an ER-like compartment **B**. Growth assay of TgGC iKD cells under 0,1,10%FBS shows TgGC cannot be rescued by excess serum. **C i**. Conoid extrusion assays using phosphodiesterase inhibitors Zaprinast and BIPPO at indicated concentrations on WT tachyzoites with 5µM A2387 as a positive control. n=8 Biological replicates, error bars represent SEM. **C ii-vi**. Conoid extrusion on various mutant cell lines and 0.5 µM Compound 1. A final concentration of 5µM final A23187 (Ca2+) or 2.5µM BIPPO (cGMP) was used for experiments on mutant cells unless otherwise indicated. For TgGC n=4, TgRNG2 n=8, TgCDPK1 n=4, C1 n=6, CDPK3 n=4 biological replicates. For all graphs, error bars represent SEM. P<0.05, \**x* <0.05, ** *x* <0.01, *** *x* <0.005, **** *x* <0.001.

TgGC-depleted cells were found to have a severe growth defect caused by an inability to secrete micronemes and egress from the host cell (Bisio et al. 2019; Yang et al. 2019). However, the effects of variable serum concentration were not explored. The presence of a phospholipid transporter ATPase domain in TgGC is still ambiguous and we hypothesized that TgGC might have roles in lipid uptake, possibly via the host cell. To test whether TgGC has a role in nutrient (specifically lipid) uptake, we grew an ATc regulated TgGC inducible knockdown mutant (Yang et al. 2019) with and without ATcin variable serum conditions to see if TgGC-depleted parasites could be rescued in a lipid rich environment, as has been shown in other mutants involved in phospholipid metabolism (Amiar et al. 2020). Interestingly, we found that this growth defect could not be rescued by growing parasites in excess serum (**FIgure 1B**).

### The cGMP signal chain regulates conoid extrusion

Microneme secretion and conoid extrusion are both critical processes that are known to occur in extracellular parasite when searching for a new host cell and subsequent invasion of that host (Katris et al. 2014). The cGMP signal pathway is well established as having a crucial role in activating microneme secretion for subsequent egress and invasion of host cells (Wiersma et al. 2004). However, any effects on the conoid upon inhibition of the cGMP signal pathway have been overlooked (Wiersma et al. 2004; Brown and Sibley 2018; Yang et al. 2019; Bisio et al. 2019). Given the apical localization of TgGC, we hypothesized that the cGMP signal network might have a role in triggering conoid extrusion in *Toxoplasma* tachyzoites. To test this, we harvested extracellular tachyzoites and stimulated them with cGMP phosphodiesterase (PDE) inhibitors Zaprinast and BIPPO, which are known to stimulate the cGMP pathway (Howard et al. 2015). We found that both Zaprinast and BIPPO were very efficient agonists of conoid extrusion (**Figure 1Ci**).

### *Toxoplasma* mutants of proteins in the cGMP signal chain are defective in conoid extrusion

Having established that the cGMP contributes to the regulation of conoid extrusion, we next asked if TgGC mutant parasites could still efficiently extrude their conoids in response to a stimulus. To test this, we treated WT and TgGC-depleted cells with either A23187 or BIPPO as before. TgGC-depleted cells extruded their conoids less efficiently in response to a stimulus, although they were relatively slightly less responsive to BIPPO than A23187 at the indicated concentrations (**Figure 1Cii**). We then observed that many proteins important for microneme secretion and invasion have yet to be investigated for defects in conoid extrusion (Brown, Long, and Sibley 2017; Lourido et al. 2010) and hypothesized that at least some known mutants might have additional conoid extrusion defects. So, we performed conoid extrusion assays using multiple parasite mutants and inhibitors and then stimulated these cells with both BIPPO and A23817. We found TgRNG2-depleted cells showed significantly less conoid extrusion upon BIPPO treatment but were mostly responsive to A23817 stimulus (**Figure 1Ciii**) similar to the response of for microneme exocytosis (Katris et al. 2014).

We next examined the involvement of Protein Kinase G, TgPKG, the primary kinase which is activated by cGMP produced by TgGC (Brown, Long, and Sibley 2017). After treatment of extracellular tachyzoites with the PKG inhibitor Compound 1, we found Compound 1-treated parasites were less able to extrude their conoids in response to either BIPPO or A23187 stimulus (**Figure 1Civ**). The same defects in both BIPPO and A23187 mediated conoid extrusion were also seen when cells were pre-treated with Compound 2, another inhibitor of TgPKG (**Figure S2A**) (Donald et al. 2006). Calcium signalling is also a well-known signalling factor responsible for conoid extrusion and microneme secretion, with a well-established model where cGMP signalling proteins TgGC and TgPKG trigger downstream intracellular calcium release to activate TgCDPK1 and TgCDPK3 for microneme secretion (Stewart et al. 2017). We thus next tested known mutants of calcium-dependent kinases involved in egress and invasion. We found that a tetracycline-inducible KO mutant of TgCDPK1 (Lourido et al. 2010) was less able to extrude its conoid in response to either BIPPO or A23187 treatment (**Figure 1Cv**). Conversely, when we examined TgCDPK3 KO cells (McCoy et al. 2012) they were able to efficiently extrude their conoids in response to both BIPPO and A23187 treatment (**Figure 1Cvi**). Taken together, these results support a previously overlooked role for cGMP signalling in conoid extrusion and that conoid extrusion is wired into the same calcium signal network regulating microneme secretion facilitating their concomitant activation for egress and invasion.

Of the proteins investigated thus far, TgRNG2 is the only protein which has not yet been investigated for calcium signalling dynamics. To examine its role, we generated a TgRNG2iKD cell line with a genetically encoded GCaMP6 calcium biosensor and tracked changes in calcium indirectly via GCaMP6 fluorescence (**Figure S2B**). TgRNG2iKD+GCaMP6 cells were pre-treated with or without ATc, stimulated with BIPPO and tracked over 60 seconds. We found that TgRNG2 depleted cells showed less BIPPO-triggered calcium release (**Figure S2B**), suggesting that TgRNG2 interacts in some way in the cGMP signal pathway controlled by TgGC. Furthermore, we found that TgRNG2 depleted cells were slower to respond to BIPPO-triggered egress (**Figure S2C**) but were responsive to A23187 (**Figure S2D**), broadly consistent with a defect in the cGMP signal pathway that can be rescued by a downstream calcium signal (Katris et al. 2014). Taken together, TgRNG2 acts within the cGMP signalling pathway and is apparently implicated in signal amplification through calcium release, although the mechanism of this role is not fully clear.

### TgGC-depleted parasites display both disrupted phospholipid metabolism and signalling

In diverse eukaryotes, both phospholipid transporter and cGMP cyclase proteins are well known to be important for lipid trafficking and lipid signalling respectively. So, we thought that the Alveolate specific GC would represent a unique opportunity to examine a novel biological example of a protein responsible for regulating lipids for both these processes simultaneously. We therefore examined changes in lipids involved in trafficking and/or signalling upon TgGC knockout by performing an exhaustive fatty acid and phospholipid analysis by Gas Chromatography Mass Spectrometry (GC-MS). The fatty acid composition of TgGC-depleted cells showed key differences in major fatty acid species (**Figure 2A**). First, there was a notable increase in the remnant plastid (apicoplast) derived C14:0, and ER derived C20:1 (Ramakrishnan et al. 2012) in TgGC-depleted cells pre-treated with ATc (**Figure 2A**). Both the apicoplast and ER are major metabolic hubs for phospholipid production (Amiar et al. 2016), suggesting that the parasites might be adapting by upregulating phospholipid production when TgGC is knocked out. There was also a notable decrease in oleic acid 18:1 in TgGC-depleted cells (**Figure 2A**).

**Figure 2:**
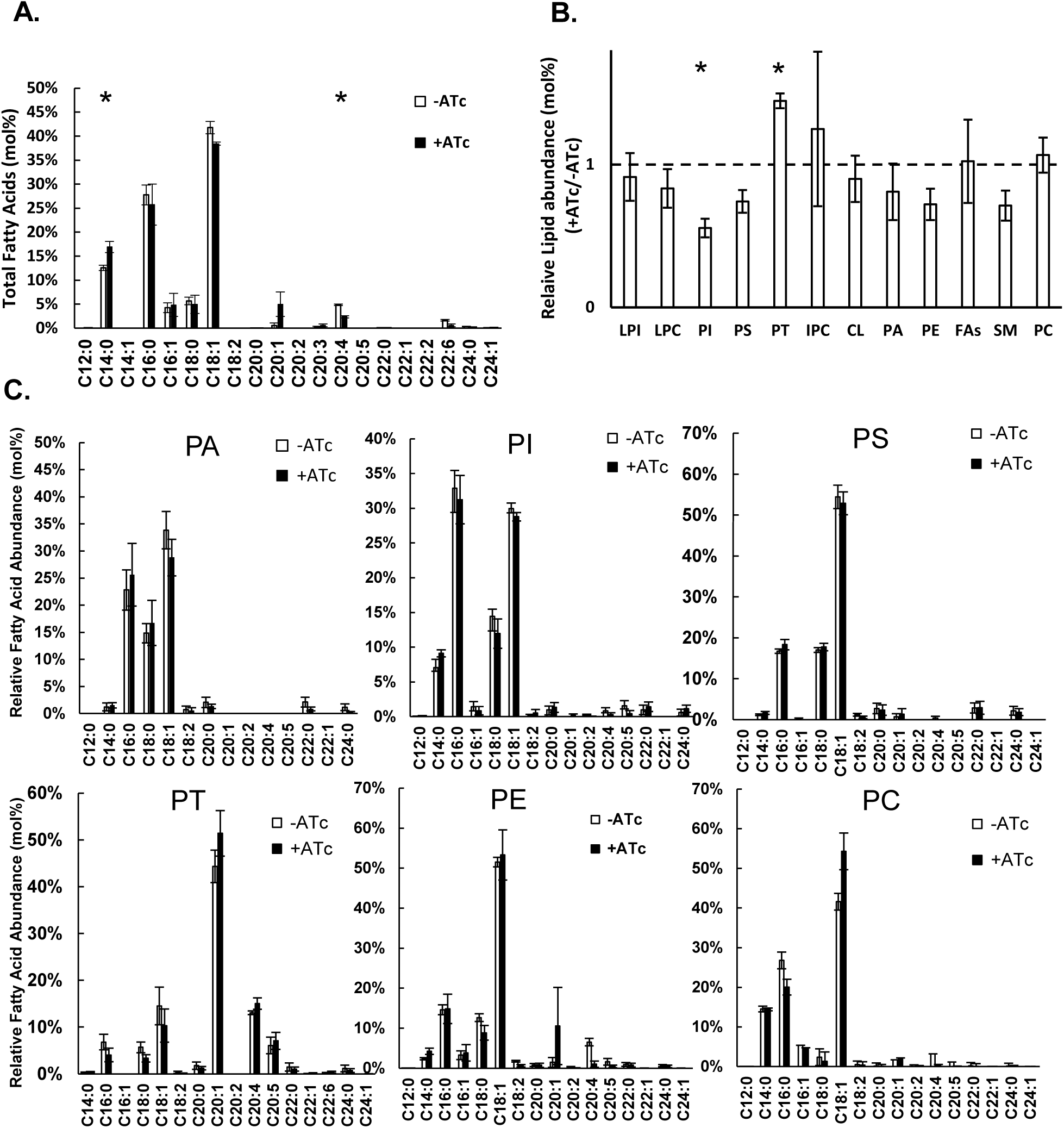
Lipidomic analysis of TgGCiKD suggests a role for TgGC in Phosphatidiylinositol signalling. **A**. Fatty acid profile(mol%) of TgGCiKD cells treated with or without Atc **B**. GC-MS analysis of lipids extracted by TLC, expressed as a ratio +ATc/-ATc **C-H**. Fatty acid profiles of PA, PI, PS, PT, PE and PC of TgGC with and without ATc pre-treatment. For all graphs, TgGC: n=3 biological replicates. P<0.05, \**x* <0.05. All error bars represent SEM.

Unsaturated fatty acids are important for increasing membrane fluidity during movement and it is possible that the reduction in 18:1 represents a decrease in membrane fluidity during extracellular motility. Additionally, C20:4 (arachidonic acid) is significantly reduced in cells with TgGC-depleted after ATc treatment (**Figure 2A**), which is interesting as lipid intermediates of the PI cycle are commonly enriched in C20:4 (Epand 2017).

When we examined individual lipid classes, we observed that TgGC-depleted cells have broadly similar levels of lyso-phosphoinositol (LPI), lyso-phosphatidylcholine (LPC), inositol phospho-ceramide (IPC), cardiolipin (CL) and free fatty acids (FAs) (**Figure 2B**). Parasites with TgGC-depleted after ATc treatment exhibited slightly reduced levels of phosphatidylserine (PS), phosphatidic acid (PA), phosphatidylethanolamine (PE) and sphingomyelin (**Figure 2B**). However, by far the most significantly reduced lipid in TgGC-depleted cells was phosphatidylinositol (**Figure 2B, S3**), which has known roles in both lipid trafficking and signalling (Shin et al. 2020). TgGC-depleted cells showed a mild increase in phosphatidylcholine (PC) abundance but the apicomplexan specific lipid phosphatidylthreonine (PT) was rather unexpectedly by far the most significantly increased phospholipid in TgGC-depleted cells (**Figure 2B**).

Despite their levels being reduced in TgGC-depleted cells, the fatty acid profile of PA, PS and PI was broadly similar, although PA and PS showed slight decreases in oleic acid 18:1, and slight concomitant increases in unsaturated C16:0 and C18:0 (**Figure 2C**). In TgGC-depleted cells PT showed an overall normal FA profile but with slightly increased levels of the signature 20:1 and 20:4 FA chains for which PT is known (**Figure 2C**) (Arroyo-Olarte et al. 2015). Interestingly, the FA profile of PC showed a large increase in 18:1 suggesting that the TgGC-depleted cells are either synthesising or scavenging more of this specific PC (**Figure 2C**). We also saw a large increase in 18:1 in the profile of SM upon TgGC depletion (**Figure S3**), so that PC and SM where the only lipids to have a large increase in 18:1 (**Figure S3**). This may reflect differences in outer and inner leaflet lipid composition of PC and SM (Lorent et al. 2020). The FA profile of PE was largely similar between TgGC-depleted and non-depleted cells, however there was an increase in C20:1 and a decrease in C20:4 consistent with the total FA analysis (**Figure 2A, C**). Taken together, in TgGC-depleted cells, there are consipuous changes in phospholipid metabolism, particularly phosphatidylinositol and phosphatidylthreonine, and to some extent in oleic acid 18:1.

### Both TgGC and TgRNG2 are involved in the apical DAG-PA signalling events

The phospholipid PA is well known to be involved in signalling events that control microneme secretion in *Toxoplasma* tachyzoites. Given changes in PA levels detected in the TgGC-depleted cells (**Figure 2B**), and the known function of PA interchanged with DAG in controlling microneme secretion (Bullen et al. 2016), we wondered whether TgGC plays a role in PA metabolism during the signalling events of invasion and egress? Particularly, we wondered what the behaviour of signalling lipids such as PA might look like in real time, over the course of activation of egress and/or subsequent invasion of a host cell which occurs in a matter of minutes. To test this, we examined levels of PA over time in response to a cGMP stimulus, BIPPO, to determine whether PA-mediated events are controlled by the cGMP signal path. Given that conoid extrusion occurs within a very rapid time frame of 30 seconds (**Figure 1C**), we considered that a time frame of up to one minute would be appropriate to examine lipid kinetics in extracellular tachyzoites. TgGCiKD cells without ATc pre-treatment were stimulated by BIPPO and a rapid increase in PA was detected within an astonishing 5 seconds of stimulus with PA levels returning to near pre-treatment levels within 60 seconds (**Figure 3A**). Cells depleted of TgGC, however, showed no significant increase in PA levels after BIPPO treatment (**Figure 3A**). This suggested that cGMP generated by TgGC is required for this fast response of PA production. We then hypothesized that other proteins known to be involved in the cGMP signal path might also show abnormal PA production with these proteins inhibited or depleted. So, we investigated cells with disrupted TgPKG and TgRNG2 to see if they produce less PA upon BIPPO stimulation. Having established that 5 seconds is enough to observe a rapid change in PA, we then focused on this time point and measured PA production in parasites within 5 seconds of BIPPO stimulus in WT cells with or without pre-treatment with PKG inhibitor Compound 2 (Donald et al. 2006). We found that Compound 2 inhibited cells had lower levels of PA at 5 seconds of BIPPO stimulation (**Figure 3B**). Similarly, we quantified PA in TgRNG2iKD cells pre-treated with or without ATc, stimulated with BIPPO, and RNG2-depleted cells showed lower levels of PA within 5 seconds of stimulus also (**Figure 3C**). These data confirm that PA is a rapidly produced product of the cGMP signal chain and that TgRNG2 and TgPKG are part of the is signal network controlled by TgGC.

**Figure 3:**
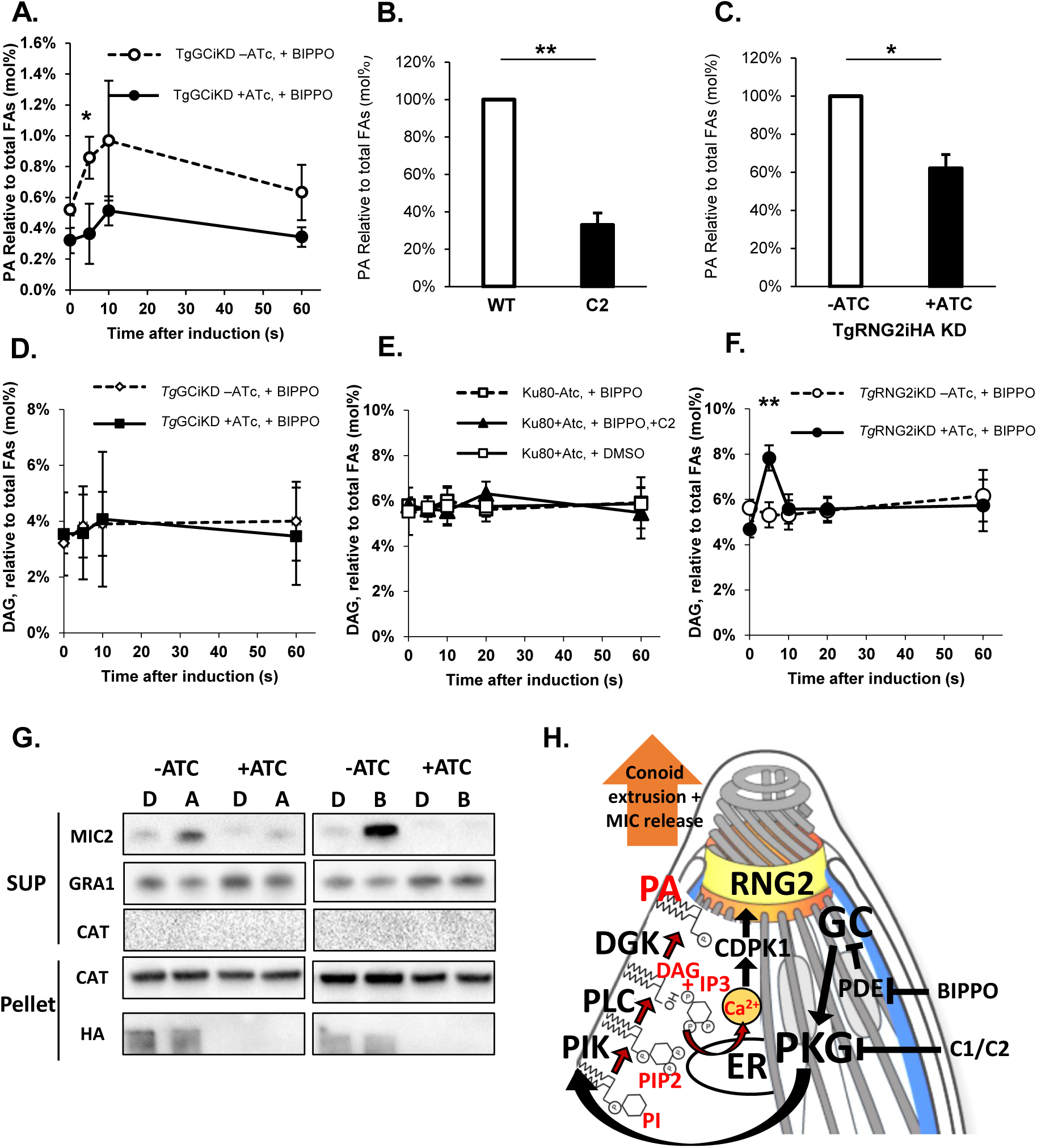
Rapid kinetics of DAG and PA are altered in TgGC and TgRNG2 mutants. **A**. Pulse experiment to analyse PA kinetics over time. TgGCiKD cells pre-treated with or without ATc were stimulated with 2µM BIPPO over 60 seconds, and PA levels (mol%) analysed at each time point. n=3 biological replicates **B**. WT tachyzoites were treated with TgPKG inhibitor Compound 2 (C2, 0.5µM) before being stimulated with BIPPO, and quenched within 5 seconds, and PA was quantified. n=3 biological replicates **C**. PA quantification in TgRNG2 cells pre-treated with ATc after being stimulated with BIPPO and quenched within 5 seconds. n=5 biological replicates **D**,**E**,**F**. Graphs showing kinetics of lipid second messenger DAG in TgGC, TgPKG and TgRNG2 inhibited cells. For TgGC and TgPKG n=3, for TgRNG2iKD n=4 biological replicates. P<0.05, \**x* <0.05, ** *x* <0.01, *** *x* <0.005, **** *x* <0.001. All error bars represent SEM. **G. S**ecretion assay of TgGCiKD cells pretreated with or without ATc, and stimulated for 20 minutes with A23187 (A) or BIPPO (B). **H**. Model for lipid second messenger turnover in *Toxoplasma* during concomitant activation of conoid extrusion and microneme secretion, beginning clockwise from TgGC.

In the known signalling events of microneme secretion DAG sits in a central location. DAG can form PA via DGK and also be restored by PA conversion back to DAG by PAP (Phosphatidic Acid Phosphatase) (Bullen et al. 2016). Furthermore, IP3 is a common signalling product of PI-PLC signalling activity. Specifically, PI-PLC dependent cleavage of PIP2, produces DAG and IP3. IP3 signal release is a common activator of Ca^2+^ store release, and in *Plasmodium* it is known to occur downstream of PKG activation (Brochet et al. 2014; Stewart et al. 2017). Therefore, DAG is at a crucial intersection, interchanging with multiple second messengers, making DAG kinetics a potentially useful readout of invasion-related signalling events. Given the fast response of PA to BIPPO-stimulation (**Figure 3A**) we sought to test if DAG changes in the same manner over time. To test this, we measured DAG kinetics over the same times as for PA. No significant change in DAG levels were seen throughout the events following BIPPO stimulus in TgGCiKD cells regardless of ATc pre-treatment (**Figure 3D**). Similarly, pre-treatment with PKG inhibitor Compound 2 also resulted in no change of DAG, nor was a change seen in parasites treated with DMSO instead of BIPPO. (**Figure 3E**). However, in TgRNG2 depleted parasites a sharp, short-lived DAG increase was seen following BIPPO stimulus (**Figure 3F**) This implies that the cGMP signal pathway initiated by TgGC is still active in TgRNG2 depleted cells and that loss of TgRNG2 is preventing the conversion of DAG to PA during microneme secretion.

TgGC has been shown to be involved in microneme secretion in *Toxoplasma* tachyzoites (Brown and Sibley 2018; Yang et al. 2019; Bisio et al. 2019). We repeated these experiments and we found that TgGC-depleted cells were unable to secrete in response to BIPPO stimulus and only partially by A23187 stimulus similar to previous reports (**Figure 3G)** (Yang et al. 2019; Bisio et al. 2019). Additionally, we tested for secretion of dense granule proteins by examining secretion of GRA1, which we found to be secreted in broadly similar amounts in both WT and TgGC-depleted cells (**Figure 3G**). Dense granule proteins are secreted independently of micronemes and therefore serve as a good indicator of both the specificity of a process and also parasite viability. Viability is also supported by the lack of CAT release into the supernatant showing that parasites are likely intact and not lysed (**Figure 3H**). Taken together, these experiments place TgGC upstream in the lipid signal chain, likely as the master regulator of lipid signalling controlling both conoid extrusion and microneme secretion in *Toxoplasma* tachyzoites (**Figure 3H**).

### Extracellular Toxoplasma tachyzoites readily take up exogenous PA as part of normal membrane remodelling processes

The P-type ATPase domain function of GC is hotly debated in both *Plasmodium* and *Toxoplasma* parasites. The ATPase domain is a predicted phospholipid transporter/flippase with some proposing that it may possibly transport calcium or K^+^ ions (Yang et al. 2019), and others even showing a role for responding to changes in plasma membrane lipid composition caused by excess phosphatidic acid (Bisio et al. 2019). We also can see from in silico analysis, contrary to previous reports (Gao et al. 2018), that the ATPase domain of TgGC is predicted fully functional with all conserved residues intact for activity of the flippase domain (**Figure S1C**), although *in vitro* experimental evidence for phospholipid transport is currently lacking.

Recently a novel class of transporters has emerged which include HsNir2 and Oxysterol Binding Proteins (OSBPs) which are important for maintaining ER-PM contact sites (Kim et al. 2015; Shin et al. 2020; Peretti et al. 2008). Notably, at least one of these transporters, HsNir2 has been shown to be important for importing PA from the outer leaflet of the plasma membrane towards the ER to fuel phospholipid synthesis. This is the important final step of the phosphatidylinositol cycle in which PA is recycled from the outer membrane to maintain phospholipid homeostasis during PIP signalling after vesicle secretion. (Kim et al. 2016; 2015). In *Toxoplasma* the closest homologue to HsNir2 is TGGT1_289570, which bears the putative phospholipid transfer domain, phosphatidic acid-binding PH domain, and additionally, an OSBP domain which in mammalian cells acts both as a lipid transporter and as a tether for ER organelle localization with proteins like HsNir2 and the cofactor HsVAPB (Mesmin et al. 2013). Many Oxysterol-related Proteins also bind lipids such as PI4P (Antonny, Bigay, and Mesmin 2018; Mesmin et al. 2013). In the *Toxoplasma* genome we can find a full set of proteins necessary to complete the recycling step of the phosphatidylinositol cycle (Figure S6). Therefore, given the existence of a complete set of putative PA transporters in *Toxoplasma*, we consider it unlikely that the ATPase domain of TgGC is dedicated to importing PA from the outer plasma membrane leaflet, although we cannot rule out the possibility that TgGC transports phosphatidic acid (and maybe even additional phospholipids) from the outer membrane.

Given the existence of a complete phosphatidylinositol cycle for recycling PA from the outer leaflet, we first wanted to test if phosphatidic acid could be internalized by extracellular parasites as part of the PI cycle. To test if PA can be imported, we exposed extracellular parasites to PA18:1, PA16:0 and control PC18:1, and measured their uptake to gauge which lipids are imported as part of membrane remodelling during extracellular stages. Given the extremely rapid time with which parasites extrude their conoids and turnover lipid second messengers, we chose 30 seconds as an incubation time for exposure to exogenous PA, before placing on ice to stop the lipid uptake. In this short time we found that PA is taken up extremely rapidly and localizes primarily to the peripheral plasma membrane and a cytosolic/endomembrane-like compartment that avoided the nucleus, occasionally localizing at internal puncta (**Figure 4A, S4A**). The localization of PA appeared to be broadly similar in parasites exposed to both exogenous PA16:0 and PA18:1. PC18:1 was taken up less readily over a short time scale, but after an hour of incubation, there was more intense fluorescence in what appeared to be the cytosol and at internal dots/puncta, likely endomembrane compartments (**Figure 4A**). Interestingly, when quantified by FACS, PA 18:1 was taken up more readily than PA16:0 suggesting that the unsaturated double bond in 18:1 was better at disrupting the plasma membrane to increase fluidity to then become internalized as part of normal membrane remodelling (**Figure 4Bi**). However, PA 16:0 was taken up more than PC18:1 suggesting a primarily head-group-dependent selection process for lipid internalization/membrane remodelling (**Figure 4Bii**). As an independent control, we also tested PA and PC uptake using a microscopy based method similar to published microscopy based quantification (Gras et al. 2019) in which parasites incubated with NBD-PA or NBD-PC were smeared onto coverslips coated with PEI and blocked overnight (**Figure S4**). An Immunofluorescence assay was then performed to label parasites with anti-SAG. The SAG1 provided an unbiased overlay to identify parasites and then parasites were measured for 488 fluorescence, indicative of NBD-lipid uptake. Using this method, we found broadly similar results to the FACS methods (**Figure S4B, C, D**). These results show that parasites actively take up phosphatidic acid as part of membrane remodelling during extracellular events.

**Figure 4:**
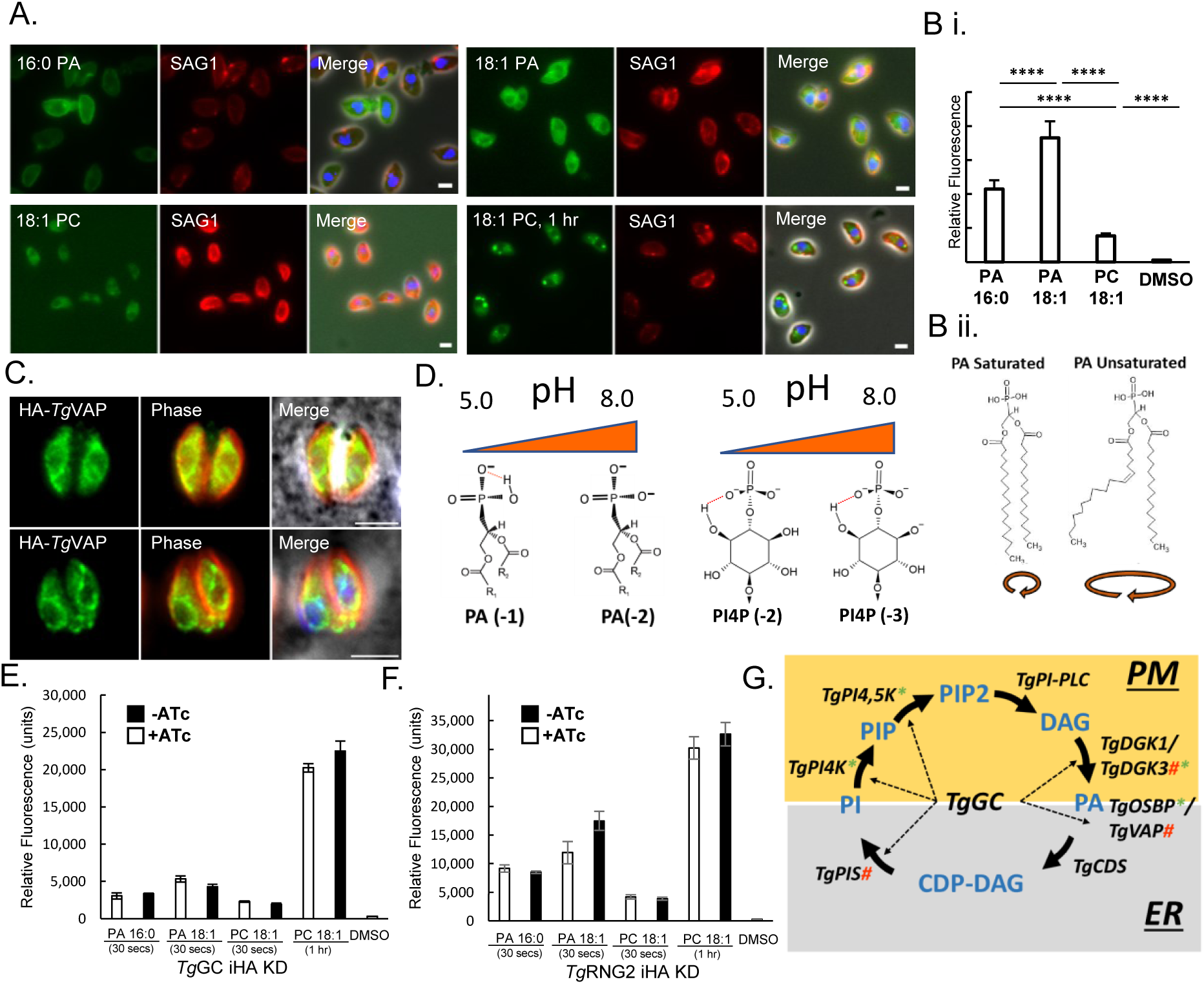
Extracellular *Toxoplasma* tachyzoites readily take up PA as part of membrane remodelling to balance the phosphatidylinositol cycle. A. Lipid uptake assay in extracellular *Toxoplasma* tachyzoites analysed by fluorescence microscopy. Extracellular parasites were incubated in DAG buffer with 5µM of indicated fluorescent NBD-lipid over 30 seconds (or 1hr where indicated). **B. i** Quantification of lipid uptake in extracellular *Toxoplasma* tachyzoites analysed by FACS comparing uptake of 5µM NBD-PA16:0, NBD-PA18:1, NBPC18:1 or equivalent DMSO control. n=24 biological replicates. ii. Structural model of saturated and unsaturated PA and the effect on lipi dbilayer fluidity due to packing of lipids. (image adapted from Avanit Polar Lipids, www.avantilipids.com) **C**. Immunofluorescence of TgVAPB by 3xHA tag. TgVAPB localizes to an ER-like localization at the periphery of the nucleus. **D**. Chemical structure indicating the variable charge state of PA and PI4P with increasing or decreasing pH (**<u>van Paridon et al. 1986, Abramson et al. 1964</u>**). **E, F**. FACS analysis of uptake of indicated NBD fluorescent lipids at indicated time points in TgGCiHA KD and TgRNG2iHA KD pre-treated with or without ATc. For TgRNG2iKD, n=4 biological replicates, and TgGCiKD n=3 biological replicates. For All graphs, error bars represent SEM.**G**. Complete model of the PI-Cycle in *Toxoplasma gondii*. Green Asterisk (*) represents protein which shows at least one altered phosphorylation site in a homologous protein in either PbGCb KO or Compound 2 treated cells in a previous Plasmdoium bergehi study (**<u>Brochet et al. 2014</u>**). Red Hashtag (#) represents gene which shows significantly altered transcription in a homologous protein of a PbGCb KO cell line in a previous Plasmodium study (**<u>Brochet et al. 2014</u>**).

We also tested phospholipid uptake in intracellular tachyzoites and we found that neither PA 16:0 or PA 18:1 could be taken up by the parasites during intracellular stages but PC18:1 clustered around the parasite vacuole and in between the parasites suggesting that PC, but not PA, could readily be taken up during intracellular host scavenging, consistent with previous reports (**Figure S4F**) (Charron and Sibley 2002). To test if similar phospholipid uptake occurs in other apicomplexan paraasites, we examined the closely related apicomplexan parasite *Plasmodium falciparum*. We incubated blood stage *P. falciparum* with either PA 16:0, PA 18:1 or PC 18:1 for 48 hours to see which lipids could be imported during replication. We found that neither PA16:0 nor PA 18:1 was taken up by the *Plasmodium* parasites during blood-stage replication (**Figure S5**), similar to *Toxoplasma*. When we exposed blood stage *Plasmodium* parasites to growth with PC 18:1 we found that fluorescent PC was readily up-taken and co-localized with DAPI indicative of dividing *P. falciparum* parasites (**Figure S5**) which is consistent with previous reports that PC is readily up-taken during blood stage growth (Grellier et al. 1991). This is in contrast to recent reports that PC cannot be scavenged by blood stage *Plasmodium* (Brancucci et al. 2017) suggesting a primarily 18:1 fatty acid chain dependent uptake compared to the PC 16:0 used in other studies (Brancucci et al. 2017). *Plasmodium falciparum* blood stage PC uptake is also consistent with what was seen here in *Toxoplasma*. We also incubated mosquito-stage sporozoites with NBD-PA and NBD-PC and found that sporozoites readily took up both PA 16:0, PA 18:1 and PC18:1 (**Figure S5**). These experiments show that phospholipid uptake as part of membrane remodelling is also important in *Plasmodium* parasite development.

To get further insight into the PA transporter family, we attempted to tag the identified family of proteins. The *Toxoplasma* genome encodes for three putative OSBPs; TgOSBP1 (TGGT1_289750), TgOSBP2 (TGGT1_264670) and TgOSBP3 (TGGT1_294740) (**Figure S6**). Both TgOSBP1 (closest homologue to PA transporter HsNir2 (Kim et al. 2015)) and TgOSBP2 (ScOsh1 homologue (Shin et al. 2020)) have putative PH domains which are known to bind PA, in support of their putative roles as PA importers, while TgOSBP3 has no predicted PH domain suggesting some other function (www.toxodb.org) (**Figure S6**). Both TgOSBP1 and TgOSBP2 have additional OSBP domains, which have known roles in ER lipid homeostasis hence our arbitrary naming of these *Toxoplasma* genes in this study. TGGT1_289570 also bears the putative phosphatidylinositol transfer domain present in HsNir2, while TGGT1_264670 lacks the PI transfer domain, and appears more like yeast ScOsh1 (Shin et al. 2020). Furthermore, the *Toxoplasma* genome also encodes for a putative cofactor TgVAP (TGGT1_318160), a homologue of HsVAP which assists HsNir1 (Kim et al. 2015), OSBPs and Ceramide Transfer proteins (CERTs) to localize at the ER (Peretti et al. 2008). We were unable to tag any of the TgOSBPs, but we could generate an N-terminally HA-tagged mutant of the HsNir2 cofactor, TgVAP. We found that HA-TgVAP was localized to the periphery of the nucleus consistent with an ER-like localization (**Figure 4C**). This is consistent with experimental evidence by hyperLOPIT in which TgVAP is assigned to the ER (Barylyuk et al. 2020), and we speculate that it could be involved in ER-PM contact sites. TgOSBP1 and TgOSBP3 are predicted to be non-essential according to a recent genome wide KO screen while TgVAPB and TgOSBP2 are have strong mutant phenotypes (Sidik et al. 2016)(www.toxodb.org) and will be the subject of future research. Taken together, these data show support the existence of a complete PI cycle in *Toxoplasma* based in the ER and PM contact sites.

Recently, TgGC was shown to be essential for secreting micronemes in response to exogenous PA. It is presumed that the exogenous PA generated instability in the plasma membrane which was then detected by the ATPase domain of TgGC and transmitted via the cyclase domain to activate motility. Furthermore, it was shown that TgGC-depleted cells were unable to egress in response to change in pH. It was thought that the phosphate headgroup of PA, with its variable charge state acts as a pH sensor by accepting protons or not depending on the extracellular pH (Young et al. 2010). However, recently PI4P has been proposed to act as a pH biosensor owing to its variable pH-dependent protonation state (Shin et al. 2020). Therefore, much of the inability of TgGC-depleted cells to respond to changes in pH could equally be attributed to PI4P or PIP2 instead of PA. Nevertheless, given that TgGC was shown to be important cells to respond to changes in PA at the plasma membrane either by membrane curvature or pH changes (Bullen and Soldati-Favre 2016), we sought to examine PA uptake in TgGC-depleted cells. We pre-treated TgGCiKD cells with or without ATc and exposed them to exogenous phospholipids and analysed by FACS. We found that TgGC-depleted cells showed reduced PA18:1 uptake over 30 seconds (**Figure 4E**). However, there was no difference in uptake of PA16:0 suggesting a chain length preference for PA uptake (**Figure 4E**). Interestingly, PC uptake was unchanged in TgGC-depleted cells over 30 seconds, but after 1 hour of incubation, TgGC-depleted cells imported slightly more PC18:1 (**Figure 4E**). Given the close proximity of TgRNG2 to TgGC, we tested phospholipid import in TgRNG2iKD cells with the hypothesis that we might see altered uptake in TgRNG2-depleted cells, which might interfere indirectly with TgGC function (**Figure 4F**). Interestingly, we found that TgRNG2 depleted cells imported more PA over 30 seconds (**Figure 4F**). Import of PA16:0 was largely unaffected after loss of TgRNG2 (**Figure 4F**). PC import was also unaffected in TgRNG2-depleted cells over 30 seconds, although there appeared to be a slight increase in PC18:1 after a 1-hour incubation, similar to TgGC (**Figure 4F**). Similar results were observed when quantified by microscopy methods (**Figure S4B**,**C**,**D**,**E**) Despite this apparent reduction in PA uptake, it is highly likely that TgGC is not importing PA, but rather regulating PA turnover as part of membrane remodelling via the TgOSBP/TgVAP complex in the PI cycle during extracellular stages, although we cannot rule out the possibility that TgGC does indeed import one or more phospholipids from the outer leaflet such as phosphatidic acid.

### Evolution of phospholipid trafficking proteins in apicomplexans and other eukaryotes

Studies on *Toxoplasma* have revealed the important role of phosphatidic acid signalling in invasion and egress. Due to its excellent genetic tractability, *Toxoplasma* has emerged in recent years as the best-studied alveolate and a model organism for investigation of the alveolate cytoskeleton. Proteins of the cytoskeleton are relatively fast evolving and therefore poorly conserved with only a handful of conserved domains identified such as alveolins (Gould et al. 2008). The emergence of phosphatidic acid as a crucial signalling lipid regulating conoid extrusion and microneme secretion at the apical complex suggest that PA regulating enzymes such as GC and DGKs could provide a framework for interpreting aspects of the evolution of parasite invasion machinery.

To illuminate the distribution and functional conservation of membrane remodelling proteins involved in PA signalling, we sought to understand their global evolution by the means of phylogenetic inference. Because the bimodular nature of apicomplexan GCs is uncommon in other eukaryotes the phylogenies of the P-type ATPase (GC:ATPase) and cyclase domains (GC:cyclase) were inferred independently (Methods). The N- and C-terminal domains in 2-domain GC:cyclase were also split and subsequently aligned together in a one-domain alignment to test whether they have co-evolved as a single evolutionary unit. Phylogenetic matrices including a broad sampling of eukaryotes were compiled by extracting homologous sequences primarily from GenBank nr and EuPathDb databases. Processed matrices were analysed by maximum likelihood tree inference.

In the Maximum Likelihood tree of 223 GC:cyclases (**Figure 5; Figure S7 S8**), the N- and C-terminal domain form two independent clades. Both clades have almost identical species presence and similar branching patterns despite the relatively short domain length (148 informative sites). Apart from the alveolates, the 2-domain GC:cyclase is present in some stramenopiles (oomycetes and a thraustochytrid) and amoebozoans. Although the N- and C-terminal GC:cyclase clades do not branch strictly as sister groups in the tree, their deep relationships are weakly resolved and their otherwise congruent evolutionary histories suggest that they once originated by tandem duplication (**Figure 5A**). The 2-domain GC:cyclase are most closely related to 1-domain GC:cyclases from eukaryotes such as humans (pdb: 4NI2_A) and further to various other proteins including the adenylyl cyclase of *Mycobacterium tuberculosis* (Guo et al. 2001). The characteristic arrangement of *Plasmodium* GC:cyclase, in which a P-type ATPase domain is fused to the N-terminus, is found in a specific subset of 2-domain GC:cyclase in alveolates and oomycetes (**Figure 5A; Figure S7 and S8**). We thus propose that the unique apicomplexan GC:cyclase architecture was present at least in the alveolate ancestor, and possibly earlier in their common ancestor with the stramenopiles (**Fig. 5C**), and originated by a single evolutionary fusion of a P-type ATPase and 2-domain GC:cyclase.

**Figure 5:**
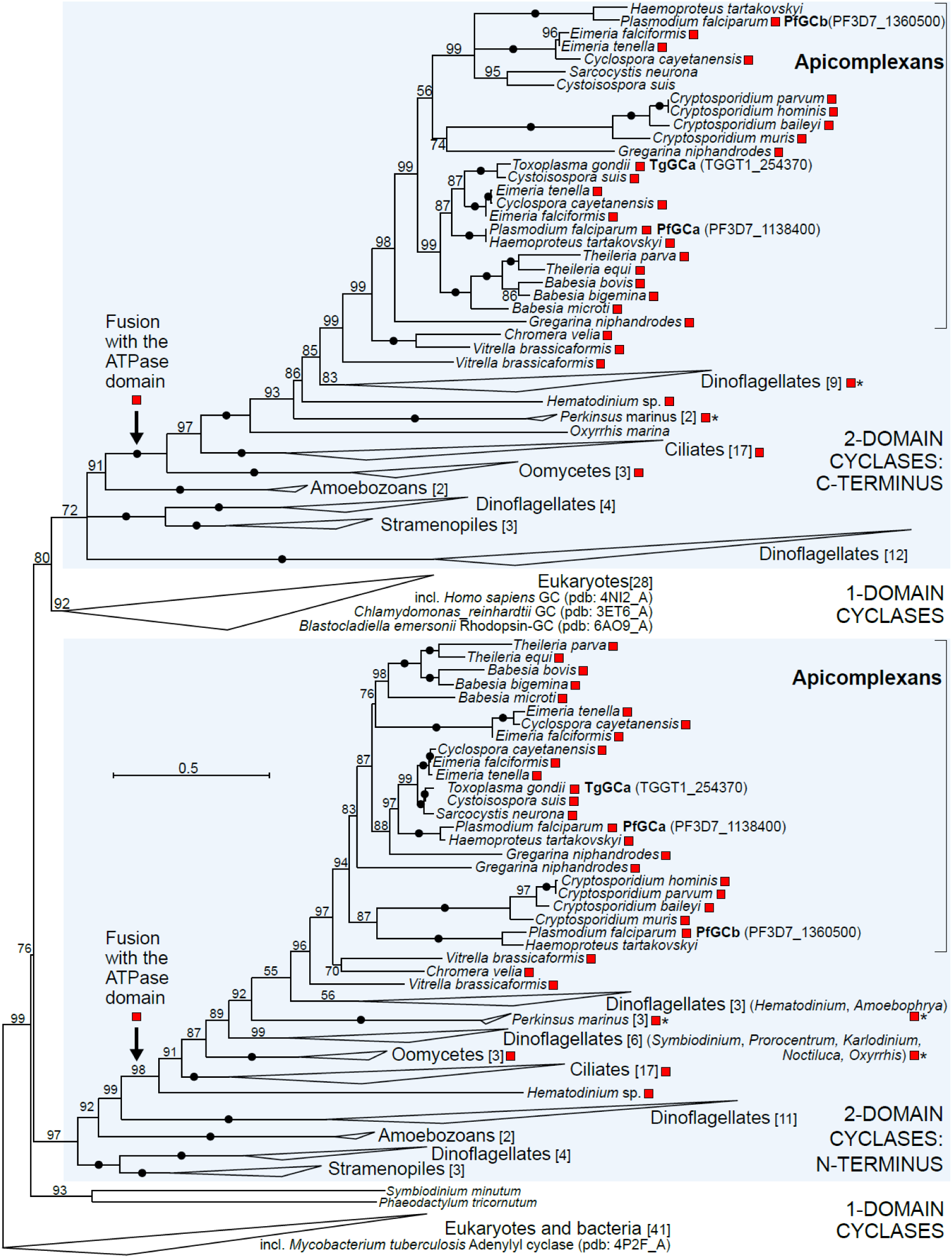
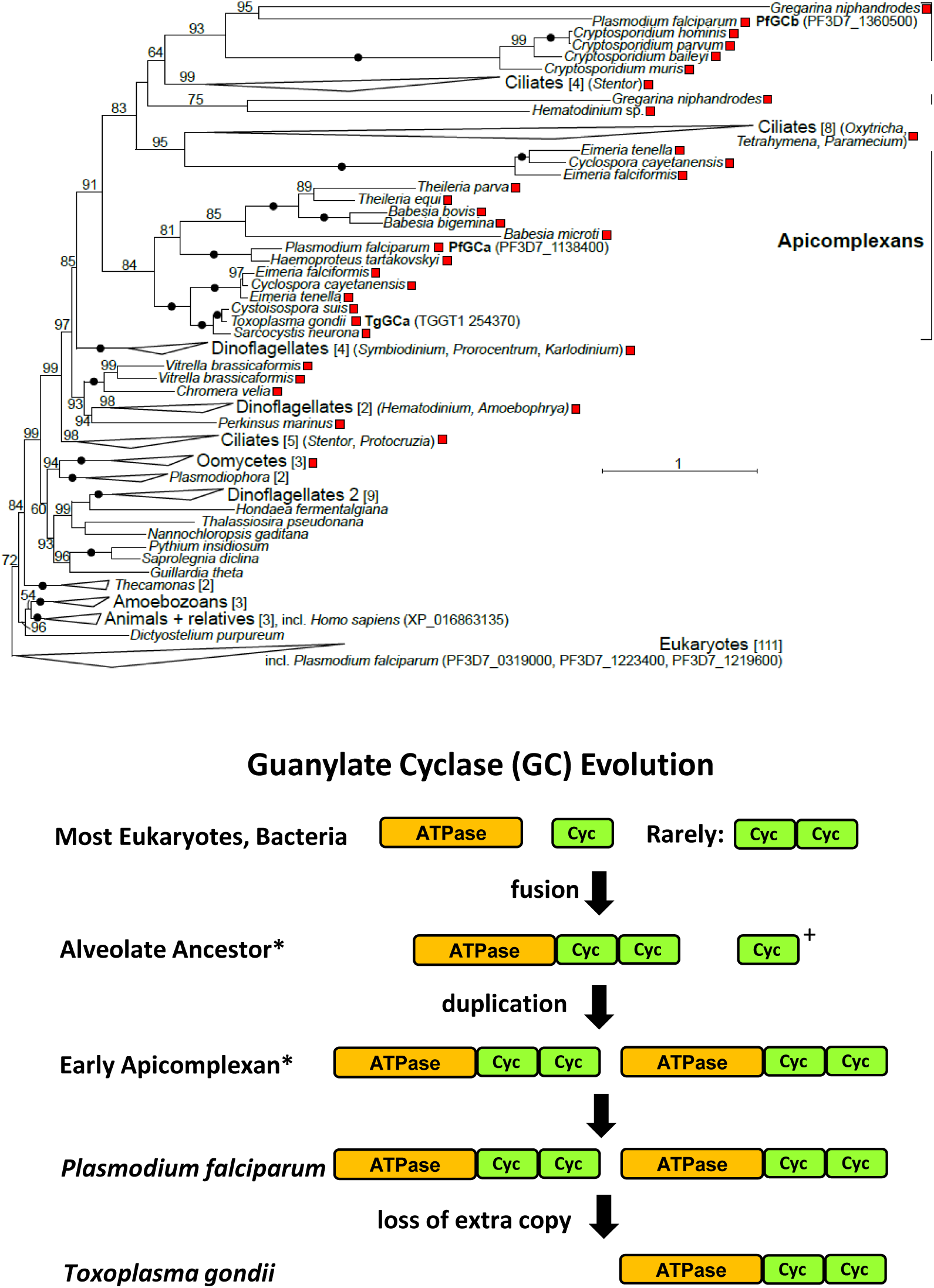
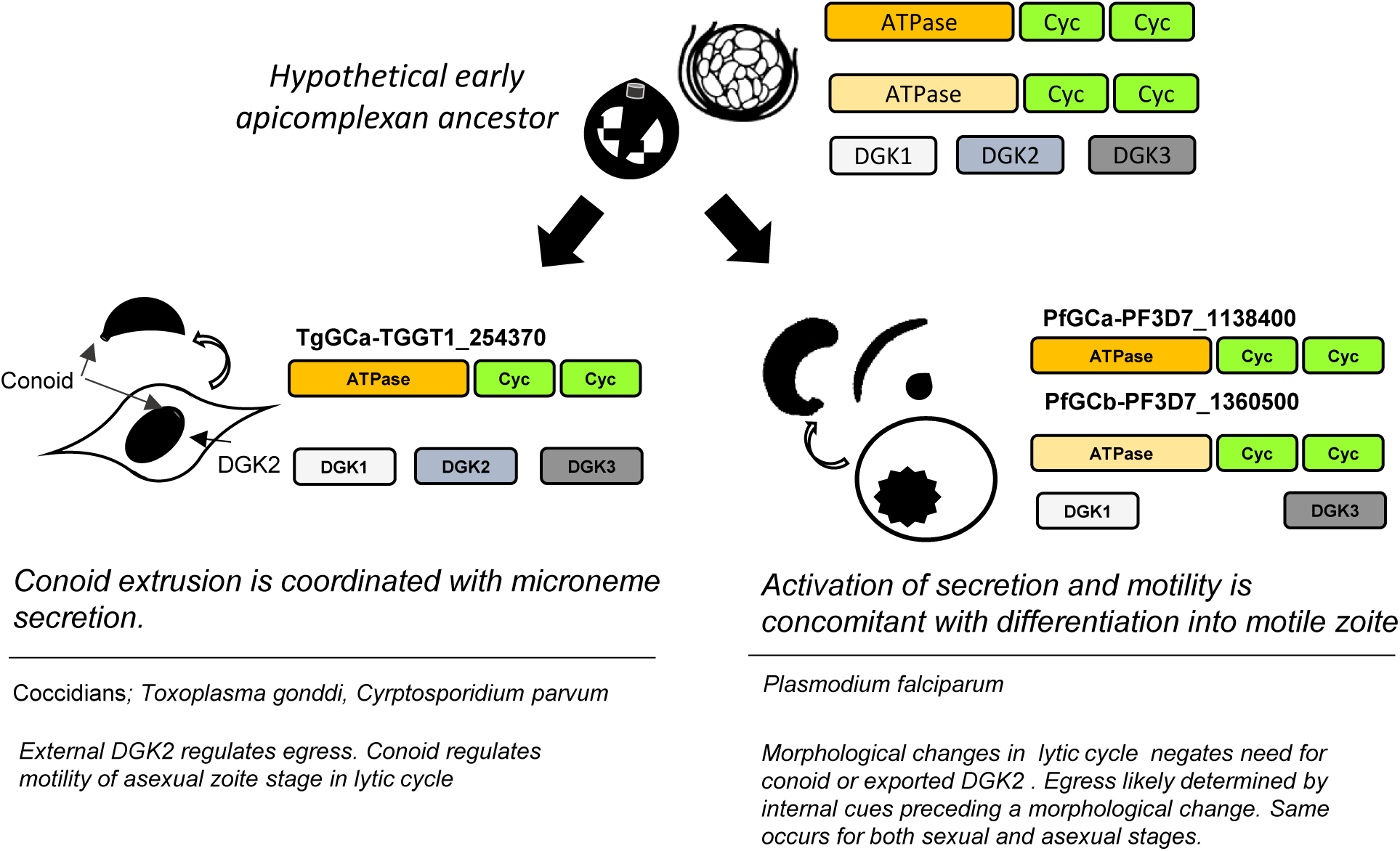
Phylogenetic analysis of phosphatidic acid signalling and trafficking proteins. Phylogenetic analysis of phosphatidic acid signalling and trafficking proteins. Phylogenetic analysis of GC in Alveolata using only the CHD cyclase domain (A) and ATPase domain (B) separately. Evolutionary scheme depicting changes in domain arrangement in GC and DGK proteins. Ancestral states represent latest predicted acquisitions of the given domain arrangement (*). Additional unfused ATPases and 1- or 2-domain cyclases were likely present in the alveolate ancestor (+).(C) Schematic of evolution of the GC protein in apicomplexan parasites. The cyclase and ATPase domains fused in the alveolate ancestor before the whole GC protein was duplicated in apicomplexans. Thesecond GC copy was lost in coccidia, but retained in Plasmodium. Toxoplasma lost the second copy of GC, but retained DGK2. Plasmodium has kept both copies of GC but lost an exported DGK2.

The phylogeny of the GC:ATPase (**Figure 5B and S9**) is more difficult to interpret, in part because many paralogs are present and evolutionary rates of individual sequences are highly variable (over 10-fold difference in branch length). The backbone of the tree is nevertheless broadly congruent with the GC:cyclase tree - all alveolate GC:ATPases group together and are placed among diverse eukaryotic ATPases that lack the GC:cyclase fusion (**Figure 5B**). The fast-evolving apicomplexan and ciliate sequences group together, in contrast to the GC:cyclase phylogeny, but this might be a long branch attraction artefact caused by strong substitutional bias. The alveolate GC:ATPase contain several paralogs (**Figure. S9**). Their relationships are not well resolved but because two variants are present in most apicomplexans (haemosporidians, *Gregarina*, and eimeriid coccidians), we conclude that an early apicomplexan underwent a GC duplication (**Figure 5C**).

Curiously, several variants of GC proteins have one of the domains either absent or substituted with another protein. The ATPase domain appears to be absent in the GCb protein of some coccidians including *Toxoplasma*, suggesting that their second copy of GC has been lost secondarily. This may be true of *Cryptosporidium* also which has only one GC variant. Notable domain substitutions include *Chromera*, which has the GC: ATPase domain fused to a domain with homology to the *Drosophila* Patched protein (Cvel_16197), and *Symbiodinium* which has GC-like proteins with an ATPase fused to a CDC50 domain, a well-known flippase co-factor (Bai et al. 2019, 50) (SmATP2-OLQ09111.1, SmATP3-OLP97098.1). Considering that these variants do not bear cyclase domains, they may not represent true GCs and may be involved in processes other than signalling. However, an assignment to GCs may be warranted by their bi-modular structure, which bears similarity to canonical GCs, and because at least the Patched domain has been shown to be involved indirectly in cAMP signalling (Lepage et al. 1995). Possibly, these alternative GCs have evolved to activate a signalling response via an atypical pathway, although it is unclear whether they remain active.

In summary, we propose that the early alveolate ancestor first fused a cyclase domain to a phospholipid transporter to couple cGMP signalling with lipid homeostasis. This dual domain GC then duplicated into at least 2x copies as can be seen in *Plasmodium*, and most other apicomplexans. The second copy of GC was then likely lost in some species such as *Toxoplasma* and *Cryptosporidium*, and is possibly in the process of loss in other coccidians such as *Eimeria* (**Figure 5C S11**).

DGKs are another group of membrane remodelling proteins known to be involved in microneme secretion and integral to the PA signalling cascade. Specifically, a parasitophorous vacuole resident protein DGK2 produces PA to activate egress via TgGC. We hypothesized that the evolutionary history of DGK2 or other DGKs might reflect changes in the evolution of GC in Alveolata. To test this, we performed a phylogenetic analysis of DGKs in alveolates to see if its evolutionary behaviour is coordinated with that of GCs. The phylogeny of DGKs (**Figure S10**) reveals a deep evolutionary divergence between the apicomplexan DGK1 and 2, and DGK3 forms. Apicomplexan DGK1 and DGK2 are each related to sequences from other alveolates and eukaryotes and were both likely present in the alveolate ancestor, having originated by an earlier duplication (**Figure S10**). Chromerids and *Cryptosporidium* contain divergent forms related to DGK1 and 2, which we call “DGK2-like”; *Chromera* also contains a distinct DGK4 form (**Figure S10)**. More detailed evolution of DGK is difficult to interpret due to the low resolution of the tree, presence of paralogs, and the loss of redundant forms: for example, DGK2 is present in coccidians but no other apicomplexans. Since DGK2 is exported to the parasitophorous vacuole in *Toxoplasma* (Bisio et al. 2019) we searched for N-terminal signal peptides in all identified DKGs by using SignalP. Although the sensitivity of this approach is limited and some DGKs have incorrectly predicted N-termini, nine sequences contained strongly predicted signal peptides. Seven of these include the DGK2 proteins in coccidians, *Perkinsus* and *Symbiodinium*, and the DGK2-like proteins in *Chromera* and *Vitrella* (**Figure S10**). It is thus possible that the chromerid DGK2-like is exported extracellularly like the coccidian DGK2 and shares similar functional aspects.

## Discussion

### Conoid extrusion is wired into the same signal network regulating microneme secretion

In summary we define the lipid signalling network controlling conoid extrusion and microneme secretion for invasion and subsequent egress in *Toxoplasma* tachyzoites and establish that it is regulated by TgGC. Conoid extrusion is a Ca^2+^ driven process and CDPK1 plays a central role in its control, but cGMP and PKG are also required for the full conoid extrusion response. Furthermore, blocking conoid extrusion by depletion of TgRNG2 reduces the efficiency of cGMP to be able to stimulate turnover of lipid second messenger DAG and downstream calcium release. This shows that conoid extrusion is wired into the cGMP and Ca^2+^ signalling pathways which also regulate microneme secretion. One possible theory is that conoid extrusion is necessary to facilitate microneme release (the retracted conoid might prevent microneme release during intracellular replication) and so their concomitant activation is coordinated by cGMP and Ca^2+^ for this reason. Recent research has found that the loss of the conoid caused by KO of TgAC9 and TgAC10 resulted in a defect in microneme secretion (Tosetti et al. 2020), consistent with the notion that conoid extrusion facilitates microneme release. While the full extent of conoid function is not known we can see that facilitating microneme release is at least one major function of the conoid and that the behaviour of conoid extrusion in *Toxoplasma* is intimately wired into the other signalling networks that are known to be important for microneme secretion and parasite motility and are therefore part of the coordinated behaviour of invasion and egress regulated by TgGC.

### TgGC shows defects in lipids required for both signalling and membrane fluidity

The behaviour of PA and DAG has important implications for both signalling and the biophysical properties of the parasite membrane. Firstly, the increase in overall PA is consistent with previous research that PA production is important for activation of microneme secretion (Bullen et al. 2016). We would also point out that a general increase in PA would serve to destabilize the plasma membrane more and thus increase fluidity for extracellular motility (Bullen and Soldati-Favre 2016), and the lack of an increase in TgGC-depleted cells shows that this is a cGMP dependent process. The lack of 18:1 in TgGC-depleted cells would also decrease membrane fluidity with increased tight inflexible packing of the bilayer, preventing proper membrane bending during motility. However, in TgGC-depleted cells by far the biggest lipid disruptions were the reduction of phosphatidylinositol for signalling and the concomitant increase in PT (**Figure 2**). We also saw a smaller decrease in PS which is consistent with previous research which showed an inverse relationship between PT and PS (Arroyo-Olarte et al. 2015). Smaller reductions could also be seen in sphingomyelin (SM) and phosphoethanolamine (PE) (**Figure 2**).

### DAG shows unusual kinetics in both wild type and mutant parasites

In kinetics experiments during activation of the early events of invasion, WT cells showed no increase in DAG, which could be because DAG is used in many other compartments of the parasitse such as storage lipids and that the relative amount used for signalling purposes are small (**Figure 3)** and difficult to detect. Alternatively, it could be that DAG is maintained at steady levels and turned over very rapidly. Indeed, this would appear to be the case as we saw a 30% increase in DAG in TgRNG2 depleted cells stimulated with BIPPO, before DAG dropped back down to basal levels within 10 seconds (**Figure 3**). This has several interesting implications. Firstly, given this behaviour with BIPPO stimulus, this suggests that the 30% increase in DAG is made exclusively by TgPLC for the purposes of microneme secretion (**Figure 3**), and the activity of TgPLC is extremely high and supports the idea that DAG is being turned over into PA as part of phospholipid metabolism. Secondly, this suggests that even though DAG levels remains flat in all the mutants, it is very likely that the DAG in WT cells is being turned over into PA at a consistent rate while in the TgRNG2 and TgGC mutants, and TgPKG inhibited cells (by treatment with Compound 2), DAG is not being turned over. Yet it is interesting that DAG is not accumulating excessively in TgRNG2iKD (**Figure 3E**), suggesting the presence of yet more phospholipid homeostasis mechanisms to prevent excess accumulation of DAG. It would be interesting then to find out what DAG is being turned into when it drops back to basal levels in the TgRNG2 depleted cells; back into CDP-DAG? cut up by a phospholipase? Thirdly, the fact that we see a large spike in DAG suggests that TgGC (**Figure 3E**) is active in TgRNG2-depleted cells and confirms that TgGC is the upstream regulator of the cGMP and calcium signal network, and that TgRNG2 is downstream in this process likely at the point of PA synthesis by one of the TgDGKs during microneme secretion, which is possibly blocked by improper conoid extrusion in the absence of TgRNG2. Similarly, the increase in PA import in TgRNG2 depleted cells can be seen as due to TgGC being active in TgRNG2 depleted cells and attempting to increase membrane remodelling for secretion and motility. More precise details on the interaction of TgRNG2 and TgGC are presently unknown, however we can clearly see that TgRNG2 is in the pathway downstream of TgGC and that the extrusion of the conoid facilitated by TgRNG2 is part of the cGMP signal chain of events.

**Figure 6:**
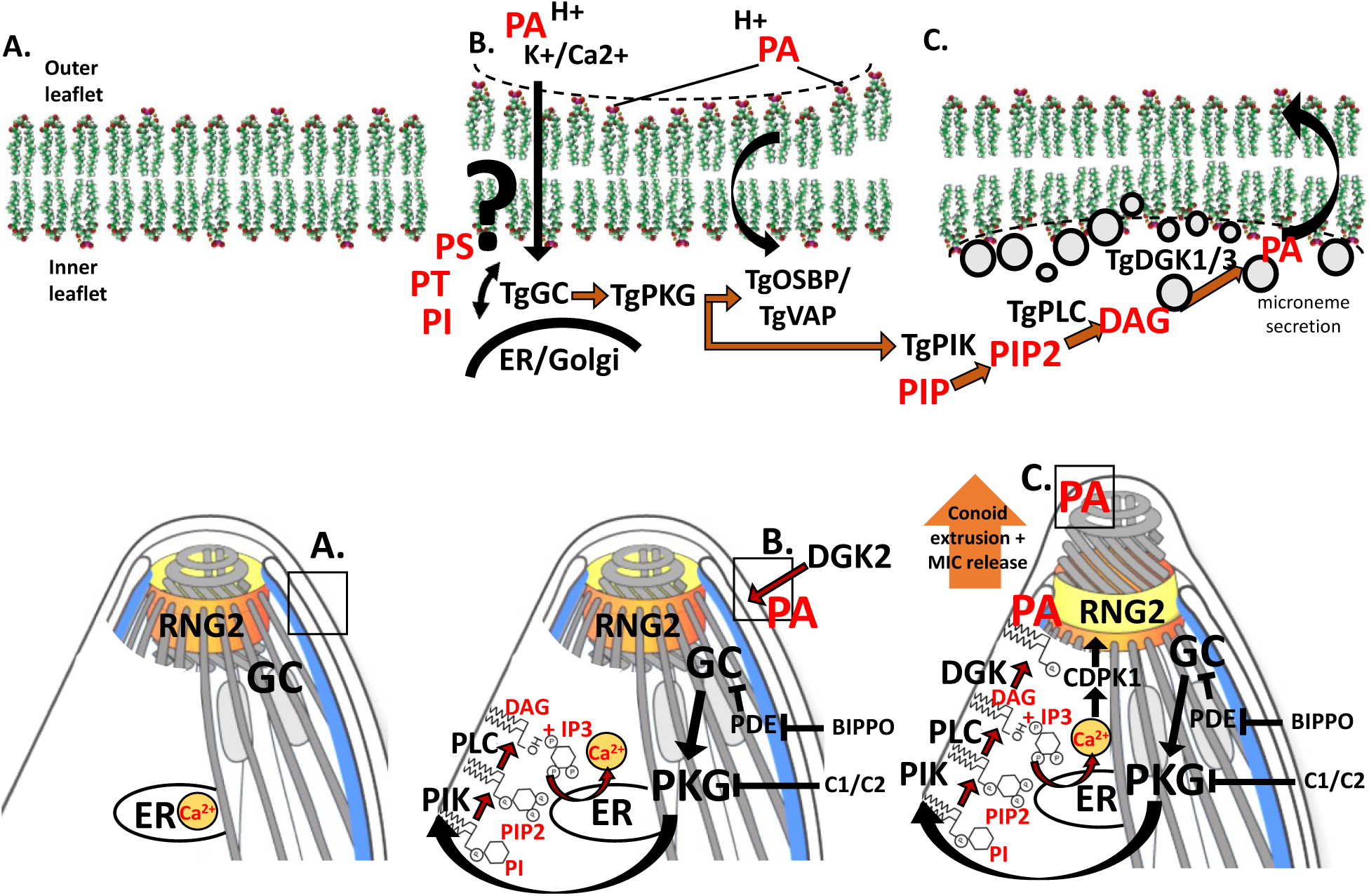
Model for PA production during cGMP and calcium signalling in *Toxoplasma*. **A**. Tachyzoite at rest during intracellular stage with stable plasma membrane **(B)** Extracellular conditions to egress result in a drop in pH at least partly attributed to exported TgDGK2 which produces PA to signal egress. Changes in the plasma membrane composition and/or pH trigger TgGC to activate PI cycle. TgGC activates TgPKG to phosphorylate PI kinase to produce PIP2 which ultimately gets cleaved to produce DAG and IP3. IP3 activates calcium release and DAG is used to produce PA at the apical complex as part of microneme secretion **(C)**. Simultaneously TgGC regulates the import of excess PA via activation of the TgOSBP/VAP complex to maintain phospholipid homeostasis **(B)**.

### The function of the ATPase domain of GC is still unknown

An unusual feature of TgGC is the presence of a P-type ATPase, phospholipid import domain, aka flippase domain, at its N-terminus. This type of accessory domain is a common feature of guanylyl cyclases in protists, whereas the mammalian GC homologue does not bear an ATPase domain. Instead there exists two isoforms of GC encoded by two separate genes; one membrane bound and one soluble/cytosolic (Johnson and Leroux 2010). Recently exogenous PA was shown to be a very potent agonist of *Toxoplasma* microneme secretion, presumably because it disrupts the plasma membrane, which triggered a secretion response regulated by TgGC. This exogenous PA mimicked an external PA source which was shown to be provided by an exported effector, TgDGK2, which was shown to be necessary for egress of *Toxoplasma* tachyzoites from the parasitophorous vacuole. However, the role of the ATPase in this PA signalling cascade was still rather mysterious and P-Type ATPase activity has yet to be shown, although predicted (**Figure S1**). Despite ours and others’ efforts, the exact function of the ATPase domain and its relationship to PA at the plasma membrane remains elusive.

### TgGC likely regulates the PI cycle

The import of PA from the outer leaflet of the PM was an important subject of research for the phosphoinositide cycle in wider eukaryotes for many years as it was unknown how end-products of phosphatidylinositol signalling were recycled to complete the PI cycle. Recently, a novel PA transporter was discovered localized to ER-PM contact sites to import PA from the external leaflet to be recycled back into CDP-DAG (Kim et al. 2015). HsNir2 is a putative PA transporter which appears to be is conserved in *Toxoplasma* and curiously has additional OSBP domains which also functions in phospholipid trafficking to and from the ER in other eukaryotes (hence our naming system of these *Toxoplasma* genes) (Shin et al. 2020; Mesmin et al. 2013) (**Figure S6**) (www.toxodb.org). Both TgOSBP1 and TgOSBP2 bear a conserved PH domain which is predicted to bind PA, but not TgOSBP3 (Kim et al. 2015; Shin et al. 2020) (**Figure S6**). Additionally, a conserved transporter cofactor homologue of HsVAPB (Kim et al. 2015) can be found in the *Toxoplasma* genome, TgVAP (**Figure 4C**), which in fibroblasts was shown to assist HsNir1 localization (Kim et al. 2015). This PA transporter HsNir2 (closest TgOSBP1 homologue), and the co-factor HsVAPB (TgVAP homologue) answered a long-standing question in wider eukaryotes regarding how PI homeostasis is maintained and how end products like PA are recycled to prevent build-up of PA at the plasma membrane and maintain phospholipid homeostasis (Kim et al. 2015). However, we cannot rule out the possibility that the ATPase domain of TgGC does indeed import PA or other phospholipids based on the limited experimental evidence presented here. Perhaps TgGC transports PI or PIPs to and from the apical cap? Although given the existence of conserved PA transporters in *Toxoplasma*, we consider it unlikely that the ATPase domain of TgGC would also specifically import PA because there already exists a conserved mechanism to recycle PA.

Interestingly, previous proteomics studies on a rodent malaria PbGCb KO cell line and Compound 2-treated cells showed that the PA transporter PbOSBP1 shows significantly altered phosphorylation of at least some sites upon disruption of the cGMP signal pathway (Brochet et al. 2014). Additionally, the putative cofactor PbVAP shows significantly reduced expression in the PbGCb KO background (Brochet et al. 2014). This evidence in *Plasmodium* suggests that the putative conserved phosphatidylinositol recycling system in *Toxoplasma* is regulated by TgGC. Given the well-established role of cGMP signalling upstream of other major calcium signalling pathways in apicomplexan parasites (Yang et al. 2019; Brochet et al. 2014), this strongly suggests that TgGC acts on TgVAP and TgOSBPs to regulate PA import as is the case for PbGCb on PbOSBPs and PbVAP. This also means that the reduced PA import seen in TgGC KO cells is much more likely to be an indirect effect of TgGC regulating membrane remodelling during active motility instead of directly importing PA which according to our proposed model, would be likely to be imported by TgOSBP1 and TgVAPB after secretion of micronemes. Similar research has shown that inhibition of endocytosis blocks motility in extracellular *Toxoplasma* tachyzoites (Gras et al. 2019). This was linked to the turnover of lipids as part of membrane remodelling through endocytosis, and that the concave lipid lysophosphatidic acid (LPA) was a key mediator of endocytosis (Gras et al. 2019). It would be interesting to examine proteomics and transcriptomics of TgOSBP or TgVAP mutants to see if a reciprocal result can be seen with expression/phosphorylation of TgGC, and this requires further research.

Altered phosphorylation of PIP kinases was a key result of cGMP signal pathway inhibition in *Plasmodium berghei* (Brochet et al. 2014), but the most striking changes were reduced expression of PbPIS, PbVAPB and PbDGK3 in the PbGCb KO, which are all proteins related to the trafficking specifically of PI or PA (Brochet et al. 2014). There was no significantly altered expression or phosphorylation of enzymes regulating DAG or CDP-DAG (Brochet et al. 2014), suggesting that regulation of phospholipids in trafficking and membrane remodelling particularly PI and PA are key features of the cGMP signal path. Interestingly, both PI and PA have variable charge state depending on the pH of the medium (van Paridon et al. 1986; Abramson et al. 1964), and have been shown to act indirectly as pH biosensors (Young et al. 2010; Shin et al. 2020), which gives some physiological understanding of why TgGC might prefer to act on these lipids and is consistent with previous observations linking pH to the role of TgGC (Bisio et al. 2019). Surprisingly, the major DAG-kinase found to be dis-regulated upon cGMP pathway ablation was PbDGK3 (PBANKA_083120) (Brochet et al. 2014), the homologue to the *Toxoplasma* TgDGK3 found in the micronemes (Bullen et al. 2016). TgDGK1 has been well-studied and found to be essential for vesicle fusion in *Toxoplasma* (Bullen et al. 2016), but the study of PbGCb KO suggests that microneme-resident DGK3, not the shorter canonical PbDGK1 homologue (PBANKA_133460), is the major DAG-Kinase responsible for microneme secretion during egress and invasion in Apicomplexa in the cGMP regulated pathway (Brochet et al. 2014). While we are still uncertain about what exactly the ATPase domain of GC does, it is clear that it is involved in regulating the PI cycle as part of a signalling process linked to sensing changes in pH. TgGC likely responds to changes in PA at the plasma membrane (Bisio et al. 2019), but based on our research it is equally likely that TgGC responds to the protonation state of PI4P or PIP2 in response to pH and the ATPase domain may even traffic PIPs to the between the apical cap and the Golgi/ER, although this theory is speculative. Taken together, according to ours and previous data, apicomplexan GC is responsible for regulating PI and PA trafficking as part of membrane turnover in the phosphoinositide cycle during extracellular motility which is likely linked to the ability of PI and PA to act as pH biosensors.

### The cGMP signal network regulates oleic acid 18:1 for membrane fluidity

Interestingly, in our results, TgGC-depletion only affected import of PA 18:1 and there was no difference seen in import of PA16:0 (**Figure 4E**,**F**). A recent study was able to differentiate the FA chain profiles of the lipids comprising the outer and inner leaflets of the plasma membrane and showed that the outer leaflet had more saturated fatty acids, while the inner membrane lipids had more unsaturated fatty acids (Lorent et al. 2020). Likely the unsaturated fatty acids incorporate into the outer leaflet and stay there, but the 18:1 is unstable in the outer layer and is transported more rapidly inside the parasite. This is probably because the 18:1 chain is less flexible and disrupts the bilayer so the parasite needs to clear this excess PA to prevent destabilizing the outer leaflet (**Figure 4B**) (Lorent et al. 2020). This shows an unexpected role for oleic acid 18:1 in increasing membrane fluidity as part of normal biophysical processes as part of extracellular motility regulated by TgGC. However, lipid uptake is seemingly largely headgroup-dependent as even PA 16:0 was imported more readily than weakly charged cylindrical PC18:1, even with an 18:1 fatty acid chain (**Figure 4**). It is presently unclear however if TgGC binds any lipids and if so, whether it has a preference for oleic acid and future research will focus on answering this question. Perhaps TgGC is able to move lipids between the apical cap and the Golgi, maybe in trafficking/signalling of phosphatidylinositol/PIPs? Interestingly, the PA examined in this study has a much higher presence of PA 18:1 than previous studies (Amiar et al. 2020) and it might be because the parasites examined here are extracellular. The increase in PA might therefore have more 18:1 in extracellular parasites and more 18:0 in intracellular parasites, and this will be an interesting topic for future research.

### Implications for conserved PA trafficking proteins in the evolution of parasitic modes of living

The identification of the dual domain Alveolate GC protein has important implications for understanding eukaryotic evolution (**Figure 5, S7-S9**), and particularly the importance of the appearance of the second copy of GC in some organisms has been under-appreciated. To our surprise, the fact that the apicomplexan ancestor had two canonical variants of GC suggests that the *P. falciparum* signal network with two GCs probably more closely resembles the apicomplexan ancestor than *Toxoplasma*, which only has one copy of GC (**Figure 5C**). Interestingly, *Eimeria falciformis* and some other coccidians have a second copy of GC which seemingly has a defunct ATPase domain, suggesting that these coccidians may be in the process of losing their second GC variant (**Figure S11**). It is interesting that GC would serve to regulate secretion of micronemes in both *Toxoplasma* and *Plasmodium*, but that *Plasmodium* would require an addtitional variant compared to *Toxoplasma*. Some have suggested a stage specific function for PfGCa and PfGCb, but this is still not clear since there is no experimental evidence for *Plasmodium* GCa in mosquito stages and *Plasmodium* GCb KO mutants have not been fully characterized in the blood stage in which they are apparently not essential (Jiang et al. 2020; Moon et al. 2009). However, given the highly conserved nature of these genes, we argue that the difference in variant number probably reflects different cellular architectures (such as apical complex differences, modes of egress etc.) and host environments in the two species (fibroblasts vs. red blood cells, etc.). It will be interesting to see how the difference in GC variant number reflects differences in coccidian and haemosporidian egress and invasion biology which could be indicative of their evolution infecting different host cells.

The phylogeny of other conserved phosphatidic acid signalling DGK proteins has also given us interesting evolutionary insights into the alveolates (**Figure S10, S11**). We find conserved TgDGK2-like homologues with signal peptides in *Vitrella* and *Chromera*. This is exciting because, to our knowledge, this is the first known effector protein exported outside of the *Toxoplasma* parasite cell which appears to have a homologue in alveolate algae. Parasite host effectors are generally poorly conserved due to the highly specific molecular nature of the parasitized host cell in question. However, it is clear that there is at least one highly conserved *Toxoplasma* dense granule protein in *Vitrella* and *Chromera* (**Figure S10, S11**), which suggests that there are probably more exported effectors that are yet to be discovered. It is presently unclear if the chromerid DGK2 is also exported, but if so it might be used to manipulate their host (e.g., if some of them exist in symbiotic fashion as suggested (Oborník et al. 2011)) or possibly to regulate their egress from a cyst stage similar to a parasitophorous vacuole (Bisio et al. 2019). Curiously, DGK2 is absent in *Plasmodium*, and this may indicate differences in its egress regulation compared to *Toxoplasma* (**Figure S11**). The major difference between the acute asexual stages of the two parasites is that *Plasmodium* asexual blood stages must undergo a developmental change into a motile zoite prior to egress in order to escape (Wang et al. 2020; Absalon et al. 2018; Rudlaff et al. 2020). It might be that *Plasmodium* has lost DGK2 because there is no requirement to signal to rings or trophozoites that have not yet developed into invasion-ready merozoites so instead *Plasmodium* possibly relies on internal metabolic cues for egress. By that same logic, perhaps elements of the conoid has also been lost or reduced in *Plasmodium* (Wall et al. 2016) because activation of egress and invasion events are more tightly linked to developmental maturation of the zoite form. In asexual *Toxoplasma* tachyzoites, there is no such delayed formation of zoites and so the tachyzoites are always invasion-ready. Thus, it is important that they have a well-regulated switch and the mobile conoid (at least in part) might control these events such as release of micronemes (Tosetti et al. 2020). Further research is needed on the full set of phosphatidic acid signalling proteins in both parasites and algae to distinguish between these speculative models.

*Toxoplasma* is an outstanding genetic model alveolate and the central role phosphatidic acid and associated proteins play in its biology has helped us identify crucial regulatory components of the alveolate cytoskeleton necessary for egress and host cell invasion. Until other genetic systems become established, *Toxoplasma* will continue to serve as a model for understudied alveolate algae and the phylum of apicomplexan parasites, which are the causative agents of numerous global diseases.

## Methods

### Parasite Culture and Transfection

*T. gondii* tachyzoites were grown by serial passage in human foreskin fibroblast (HFF) cells in DMEM supplemented with 1% Foetal Bovine Serum (FBS) as previously described (Striepen B, 2007).

### Plasmids and cloning

TgGCiKD was obtained from Yang et al. 2019 and used primarily throughout this study. TgGCiKD was also independently constructed using primers in Table S1 and inserted into equivalent cut sites of plasmid pPR2-HA3 (Katris et al. 2014). To localize TGGT1_224190 (TgFLP2), the following primers **<u>TACTTCCAATCCAATTTAGC</u>**CTCGAGTA-CTCGGTCTTGACTACG and **<u>TCCTCCACTTCCAATTTTAGC</u>**CGGCGACGCGAGCT were used to amplify the gDNA. For TGGT1_245510 (TgFLP5) primers **<u>TACTTCCAATCCAATTTAGC</u>**CAACCGGAAGGTCGTCAAGG and **<u>TCCTCCACTTCCAATTTTAGC</u>**CGCATCGTTGGCTGCGTTC were used to amplify the gDNA. Both PCR products were inserted into plasmid pLIC-HA-tubCAT. The plasmids for TgFLP2 and TgFLP5 were linearized with BglII and PstI respectively as previously described (Sheiner et al. 2011).

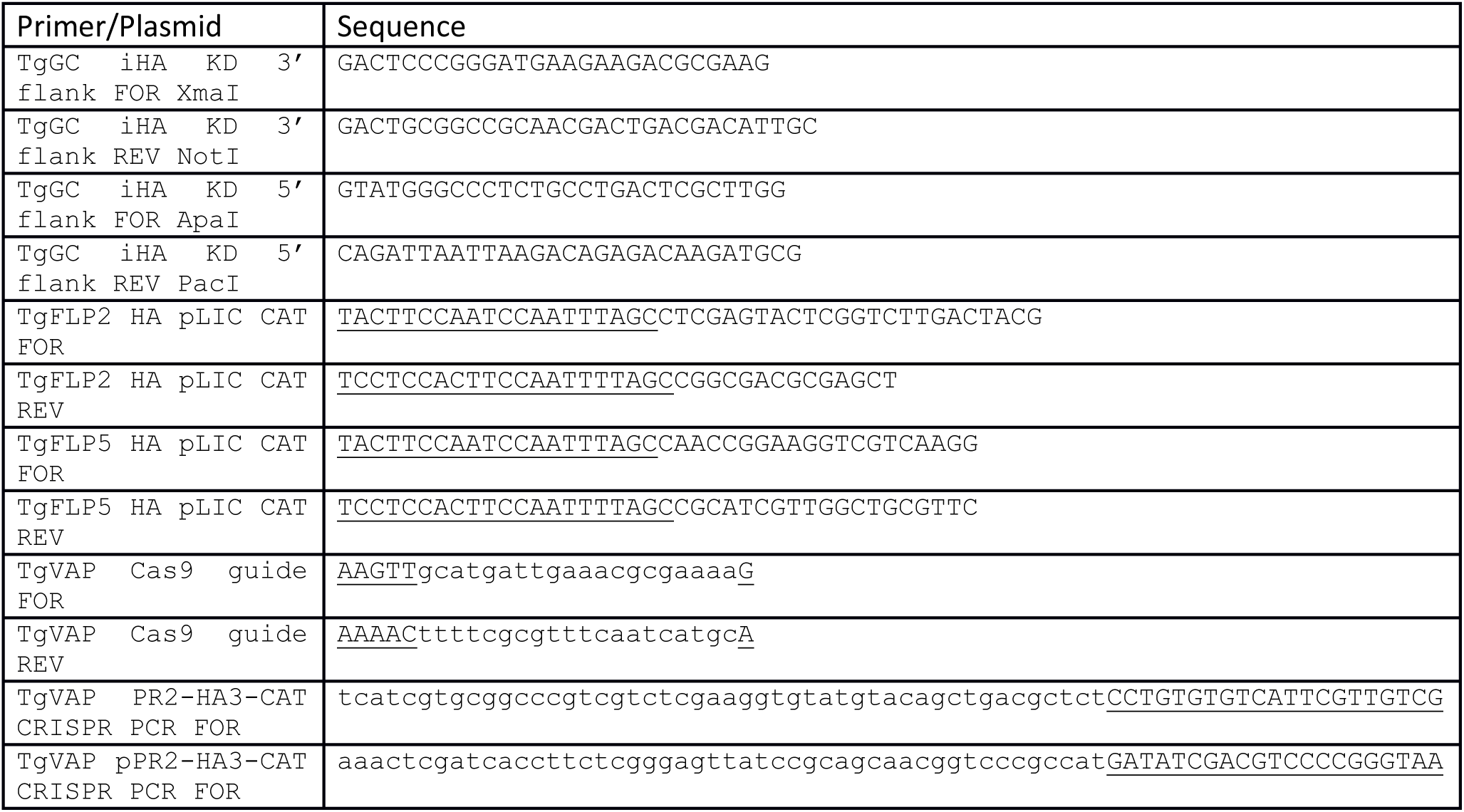

### Conoid extrusion assays

Extracellular parasites were filtered, counted and spun at 1000 xg for 10 minutes. Parasites were resuspended in DMEM (without additional supplements), to a concentration of approximately 5.0×107 parasites/ml. 100 microliters of parasites were then mixed with equivalent volumes of solutions containing double concentrations of 10 μM calcium ionophore A23187 or 5μM BIPPO to make a 200 μl solution (final concentration 5μM A23187 and 2.5μM BIPPO), or equivalent concentration of DMSO control. Parasites with agonist or DMSO were allowed to extrude conoids by shifting to a 37 °C water bath. Cells were incubated for 30 seconds and then fixed by addition of 10 μl of 25% glutaraldehyde (final concentration 1.25%), for 30 minutes. Parasites were spun at 800 rpm for 2 minutes, and resuspended in 200 μl 1xPBS. 50 microliters of parasites was then smeared onto Polyethylenimine(PEI)-coated coverslips and parasites were allowed to settle for 20 minutes, before being rinsed in deionized water and mounted onto microscope slides using 10 microliters of Fluorogel®. Parasite conoids were scored blindly by phase contrast on a 63x or 100x objective.

### DAG Pulse /Lipid kinetics

Parasites were grown for three days with or without ATc and harvested fresh on day of egress. Fully lysed cultures were filtered through a 3.0um polycarbonate filter and spun down at 1800rpm for 10 minutes. Parasites were aspirated, washed with DMEM and flasks of each treatment were pooled together, and spun down as before. Parasites were aspirated and resuspended to a concentration of 4×10^8 cells per ml and 0.5ml was each aliquoted into 1.5ml tubes and maintained at room temperature prior to treatment (Where appropriate, Compound 2 was added at this point and allowed to equilibrate for 5 minutes prior to stimulus.). Pairs of tubes with or without ATC/Compound 2 treatment were shifted to a 37 °C heat block and allowed to equilibrate for 60 seconds before addition of 0.5ml of preheated 2x DMEM solution with 5uM of BIPPO (2.5uM final) or DMSO (0 seconds). Cells were incubated for the indicated time points before quenching immediately on dry ice/ethanol for 5 seconds and placed on ice. Parasites were then spun down at 8000 rpm for 2 minutes at 4 °C, aspirated, washed with 1ml of ice cold 1xPBS, and spun down as before. Cells were washed with 1xPBS once more and spun down as before. Pellets were then placed at -80 for storage prior to lipid extraction and GC-MS analysis.

### Lipid analysis and PA quantification

Total lipids were extracted using chloroform:methanol, 1:2 (v/v) and chloroform:methanol, 2:1 (v/v) in the presence of 0.1MHCl with periodic sonication. Pooled organic phase was subjected to biphasic separation by adding 0.1 MHCl. The organic phase was dried under N2 gas and dissolved in 1-butanol. Total lipid was then separated by 2D-HPTLC spiked with 1 μg PA(C17:0/C17:0) (Avanti Polar lipids) using chloroform/methanol/28% NH4OH, 60:35:8 (v/v) as the 1st dimension solvent system and chloroform/acetone/methanol/acetic acid/water, 50:20:10:13:5 (v/v) as the 2nd dimension solvent system [72]. All spots including PA were visualized with primulin and scraped. Lipids were then prepared for GC-MS analysis in hexane (Agilent 5977A-7890B) after derivatization by methanolysis using 0.5M HCl in methanol incubated at 85°C for 3.5 hrs. Fatty acid methyl esters were identified by their mass spectrum and retention time compared to authentic standards. Individual lipid classes were normalized according to the parasite cell number and internal standard.

### Fluorescent Lipid import assay

Extracellular parasites were filtered, counted and spun at 1800rpm for 10 minutes. Parasites were washed once with DMEM and spun down as before. Parasites were resuspended in DMEM w 10mM HEPES (which we found crucial for the experiment), to a concentration of approximately 5×10^7^ parasites/ml. 100 μl of parasites were then shifted to 37°C for 1 minute to equilibrate. Parasites were then mixed with 100 μl of pre-warmed serum free DMEM w 10mM HEPES containing double concentrations of NBD-PA and NBD PC (10μM ^-1^) to make a final reaction volume of 200μl (5μM final concentration of NBD-PA or NBD-PC) Parasites were incubated for a further 30 seconds for NBD-PA, NBD-PC or 60 minutes for NBD-PC. Parasites were then quenched on ice for 2 minutes, spun down 8000rpm for 2 minutes at 4°C and aspirated. Cell pellets were resuspended in 150μL 1xPBS and fixed by addition of 80 μL of 10% Paraformaldehyde (approximately 3% final PFA) for 15 minutes at room temperature. Cells were spun down as before and resuspended in 200μl 1xPBS before smearing onto PEI coated coverslips and blocked in 2% FBS overnight at 4°C. Parasites were then probed by IFA using mouse anti-SAG1 (Abcam) and Alexafluor goat anti-mouse-546 and stained with DAPI. Slides were imaged on a Zeiss Apotome epifluorescence microscope. Phospholipid uptake was quantified with ImageJ, where the SAG1 signal was used to identify and label parasites creating an unbiased overlay. Identified parasites were then measured for fluorescent lipid uptake in the 488 channel. Alternatively, after fixation, parasites were resuspended in 1xPBS, diluted, and analysed directly by FACS for intensity of NBD uptake under 488 wavelength (Stocks of PA or PC were made up in DMSO to a concentration of 1mM, and stored at -20 °C in aliquots).

### Intracellular lipid uptake

For intracellular parasites, host cells were inoculated with tachyzoites in growth medium containing NBD-PA or NBD-PC and grown for a further 24 hours before fixation with 2.5% PFA in 1xPBS. Samples were mounted directly onto slides with fluorogel.

### GCaMP6 experiments

A construct containing the GCaMP6 plasmid was transfected into the TgRNG2iKD cell line using FUDR/UPRT selection as previously described (Stewart et al. 2017). TgRNG2iKD cells were then pre-treated with or without ATc as previously described (Yang et al. 2019). Briefly, upon harvest, TgRNG2iKD cells with or without ATc pre-treatment were filtered, spun down and resuspended in phenol-red free DMEM. Parasite were allowed to equilibrate at room temperature before being fed into a FACS and imaged in real time. Incremental concentrations of BIPPO was added to the parasites and the GCaMP6 fluorescence was detected over time. Microscopy of RNG2iKD cells was performed as previously described (Stewart et al. 2017).

### *Plasmodium falciparum* IFAs

*Plasmodium falciparum* was cultured as previously described (Botte et al., 2013). Briefly, cultures were grown in 2% haematocrit with RPMI supplemented with 10% Albumax, hypoxanthine and gentamicin. Cultures in airtight Perspex boxes were gassed with Beta-Mix gas. Parasites were grown for 24 hours in normal conditions before addition of 5 μg ml^-1^ fluorescent 16:0 PA, 18:1 PA or 18:1 PC into the culture and grown for a further 24 hours. Parasites were then harvested and smeared onto PEI coated coverslips before staining with DAPI and mounting onto slides with fluorogel.

### Mosquito stage IFAs

Live *P. falciparum* sprorozoites were incubated at room temperature using the indicated concentrations of fluorescent lipids and then mounted on coverslips and observed by confocal microscopy.

### Phylogenetic analysis

#### Assembly of phylogenetic datasets

*Plasmodium* and *Toxoplasma* GC and DGK proteins were used to identify a small set of homologous sequences in other alveolates. Of these, the proteins in *Chromera velia* (2 GCs and 4 DGKs) had the slowest evolutionary rates (as evaluated by branch lenghts in preliminary trees) and were used as representative queries in BLASTP searches against sequence databases. Sequences were retrieved from GenBank nr (a minimum of 250 closest hits for each query with an additional retrieval of selected species), GenBank pdb (all closest relevant hits; highlighted in tree figures), EuPathDb (proteins predicted from representative apicomplexan genomes) and the Marine Microbial Eukaryote Transcriptome Sequencing Project (proteins predicted from representative dinoflagellate transcriptomes). The GC was aligned in MAFFT v7.402 (Katoh and Standley, 2013) by using the --linsi algorithm and the N-terminal ATPase domain was separated (where present) from the cyclase domain region in the intervening spacer. Domains in all proteins were identified by NCBI Batch CD-search. The two cyclase domains of the *Chromera* GCa were then used as queries for another round of BLASTP searches in sequence databases as above. This was to ensure that all proteins related to the comparatively more divergent cyclase retion are retrieved (if missed in the first round of searches). The cyclase dataset, combining specific hits to “CHD”, “CYCc” or “GC” domains and generic hits to “NCS” superfamily (as determined by CD Search), was then realigned in MAFFT --linsi. The N- and C-terminal domains in 2-domain cyclases were split in the spacer region and combined together with 1-domain cyclases into a single alignment. Redundant sequences and other domain types if present (rare) were removed from all three datasets, which were then subjected to progressive rounds of streamlining. In each round, a quick Maximum Likelihood phylogeny was built in FastTree v2.1.10 (Price et al., 2010) and sequences that either belonged to distantly related protein families, were too fast-evolving, were heavily fragmented or belonged to closely related species or strains were removed.

#### Phylogenetic inference and signal peptide prediction

The three completed datasets were realigned by using the --linsi algorithm in MAFFT. Hypervariable sequence regions were trimmed in BMGE v1.1 (Criscuolo and Gribaldo, 2010) by using the -b 3 and –g 0.4 parameters. The final phylogenetic matrices of GC:cyclase (233 sequences, 148 sites), GC:ATPase (191 sequences, 569 sites) and DGK (149 sequences, 169 sites) were analyzed in IQ-TREE v1.6.5 (Nguyen et al., 2015). IQ-TREE was run by using the LG+F+I+G4 model and 1000 ultrafast bootstrap replicates to compute the branch supports (we avoided using standard non-parametric bootstraps, which are no suitable because of the short lengths of the GC:cyclase and DGK matrices). The presence of putative signal peptides in DGKs was predicted in all sequences in SignalP v4.1 by using the -euk model and default settings (Petersen et al., 2011).

## Abbreviations

FA: Fatty Acid
FFAs: Free Fatty Acids
PA: Phosphatidic acid
PI (P): Phosphatidylinositol (Phosphate)
PS: Phosphatidylserine
PT: Phosphatidylthreonine
ATc: Anhydrotetracycline
GC: Guanylate cyclase
DAG: Diacylglycerol
IP3: Inositol triphosphate
TLCs: Thin Layer Chromatography
GCMS: Gas Chromatography Mass Spectrometry.
cGMP: Cyclic guanosine monophosphate
cAMP: Cyclic adenosine monophosphate
OSBP: Oxysterol Binding Protein

## Author Contributions

**Conceptualization:** NJK, RFW, CYB

**Data curation:** NJK, YYB, JJ, CYB

**Formal analysis:** NJK, YYB, JJ, CYB

**Funding acquisition:** GIM, RFW, MFCD, CYB

**Investigation:** NJK, YYB, JJ, CSA, ASPY, AV, RJS, CJT

**Methodology:** NJK, YYB, JJ, RFW, CYB.

**Project administration:** NJK, RFW, MFCD, CYB

**Resources:** RFW, MFCD, CYB

**Supervision:** NJK, RS, GIM, CJT, RFW, CYB

**Validation:** NJK, YYB, JJ, CSA, ASPY, AV, RJS

**Visualization:** NJK, RFW, CYB

**Writing – original draft:** NJK

**Writing – review & editing:** NJK, JJ, RFW, MFCD, CYB

## Acknowledgements

We would like to thank David Sibley, Sebastian Lourido and Dominique Soldati-Favre for gifting antibodies and cell lines used in this study. We thank Oliver Billker for gifting Compound 2. This work and N.J.K, C.Y.B., Y.Y.-B, M.F.C.D are supported by Agence Nationale de la Recherche, France (grant ANR-12-PDOC-0028, Project Apicolipid), the Atip-Avenir and Finovi programs (CNRS-INSERM-Finovi Atip-Avenir Apicolipid projects), and Laboratoire d’Excellence Parafrap, France (grant ANR-11-LABX-0024). C.Y.B, C.J.T and G.I.M. are supported by the LIA-IRP CNRS Program (Apicolipid project). Lipidomic and fluxomic analysis was conducted in our GEMELI Lipidomics platform, which is supported by Region Auvergne Rhone-Alpes (IRICE GEMELI Grant) and Universite Grenoble Alpes.

---

**Figure S1:**
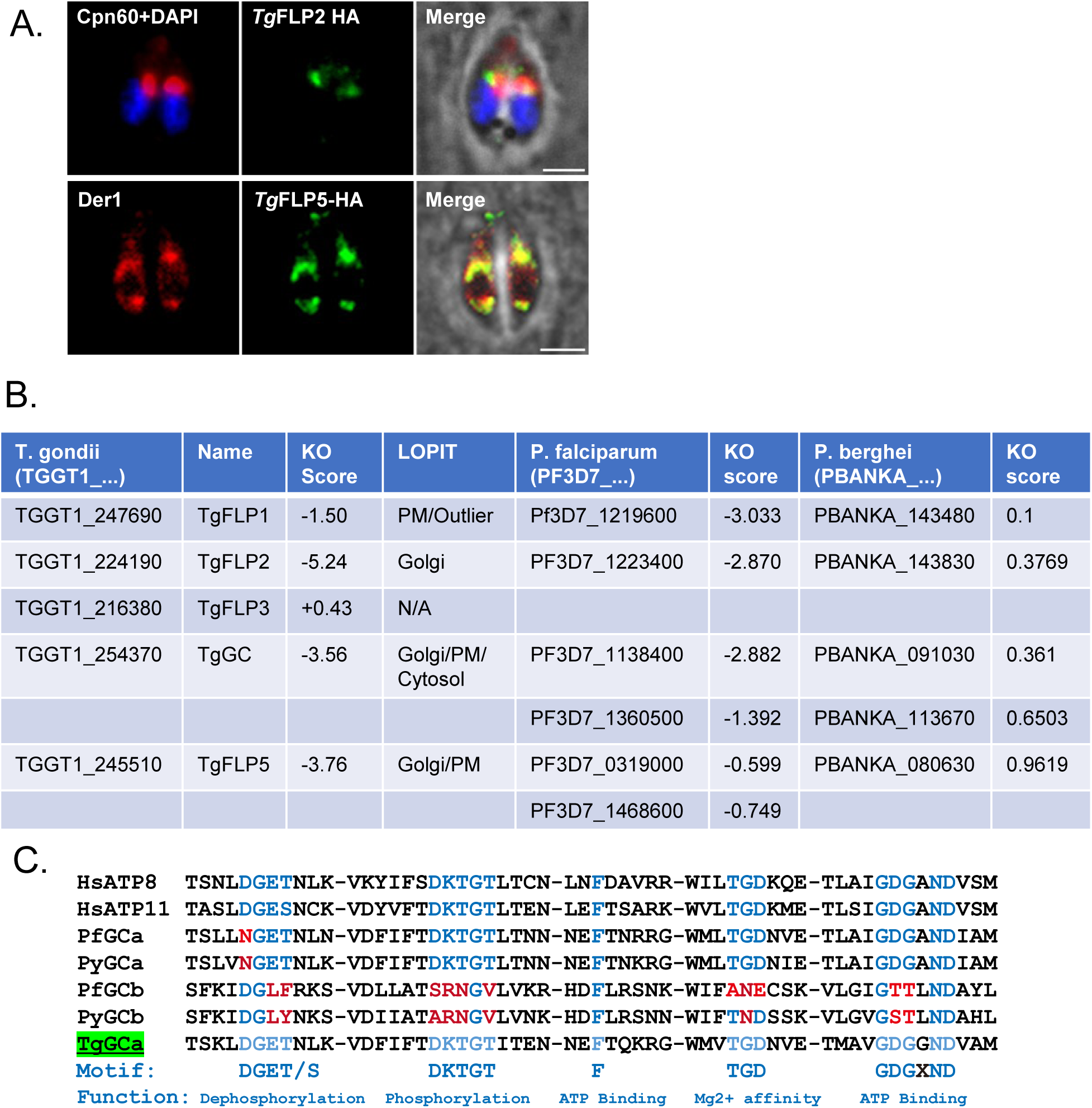
Phylogeny of phospholipid flippases in *Toxoplasma* and *Plasmodium*.. **A**. Localization of TgFLP2 (TGGT1_224190) to a Golgi–like compartment that does not co-localize with the apicoplast anti-Cpn60. Localization of TgFLP5 (TGGT1_245510) to the ER (Der1). Scale bars represent 2µm. **B**. Table of known flippases in Toxoplasma and Plasmodium with predicted localization by Hyper-LOPIT and predicted essentiality. **C**. MUSCLE Alignment of the key residues of the Human flippase ATP8 and ATP11 with the ATPase domain of Apicomplexan dual domain GCs.

**Figure S2:**
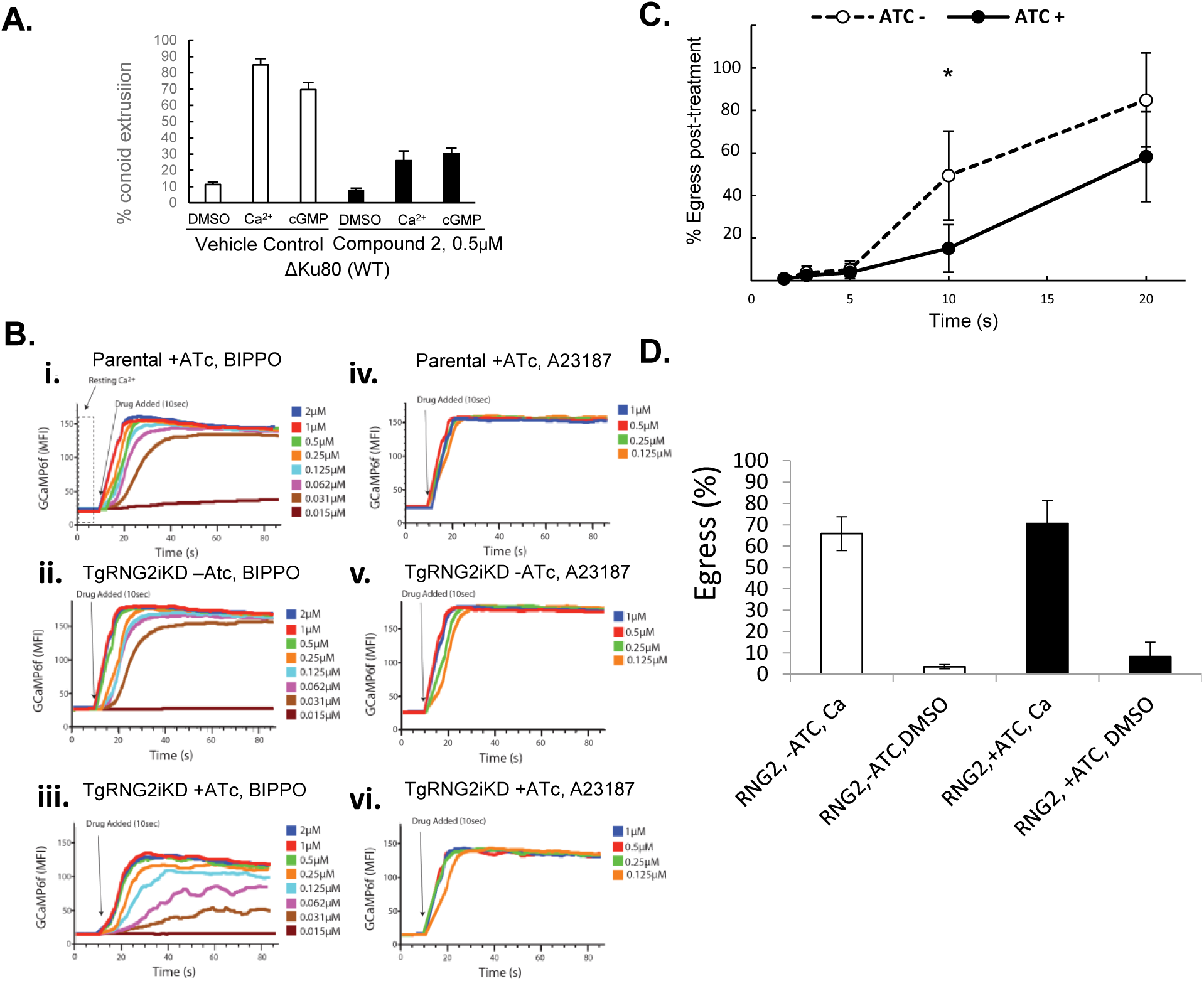
the role of cGMP signalling in conoid extrusion. **A i.** Conoid extrusion assay with WT cells and cells pre-treated with the PKG inhibitor Compound 2 (0.5uM). n=4. **B**. FACS analysis of Calcium flux in TgRNG2iKD cells expressing the calcium biosensor GCaMP6 after stimulus with either A23187 or BIPPO over time at indicated concentrations. **C**. Egress assay of TgRNG2iKD cells with or without ATc pre-treatment following stimulus with 10µM BIPPO over time. n=3 biological replicates. D. Egress assay of TgRNG2iKD cells with or without ATc pre-treatment following stimulus with 5uM A23187 after 5 minutes. n=4. For all graphs, error bars represent SEM. For all graphs P<0.05, \**x* <0.05, ** *x* <0.01, *** *x* <0.005, **** *x* <0.001.

**Figure S3:**
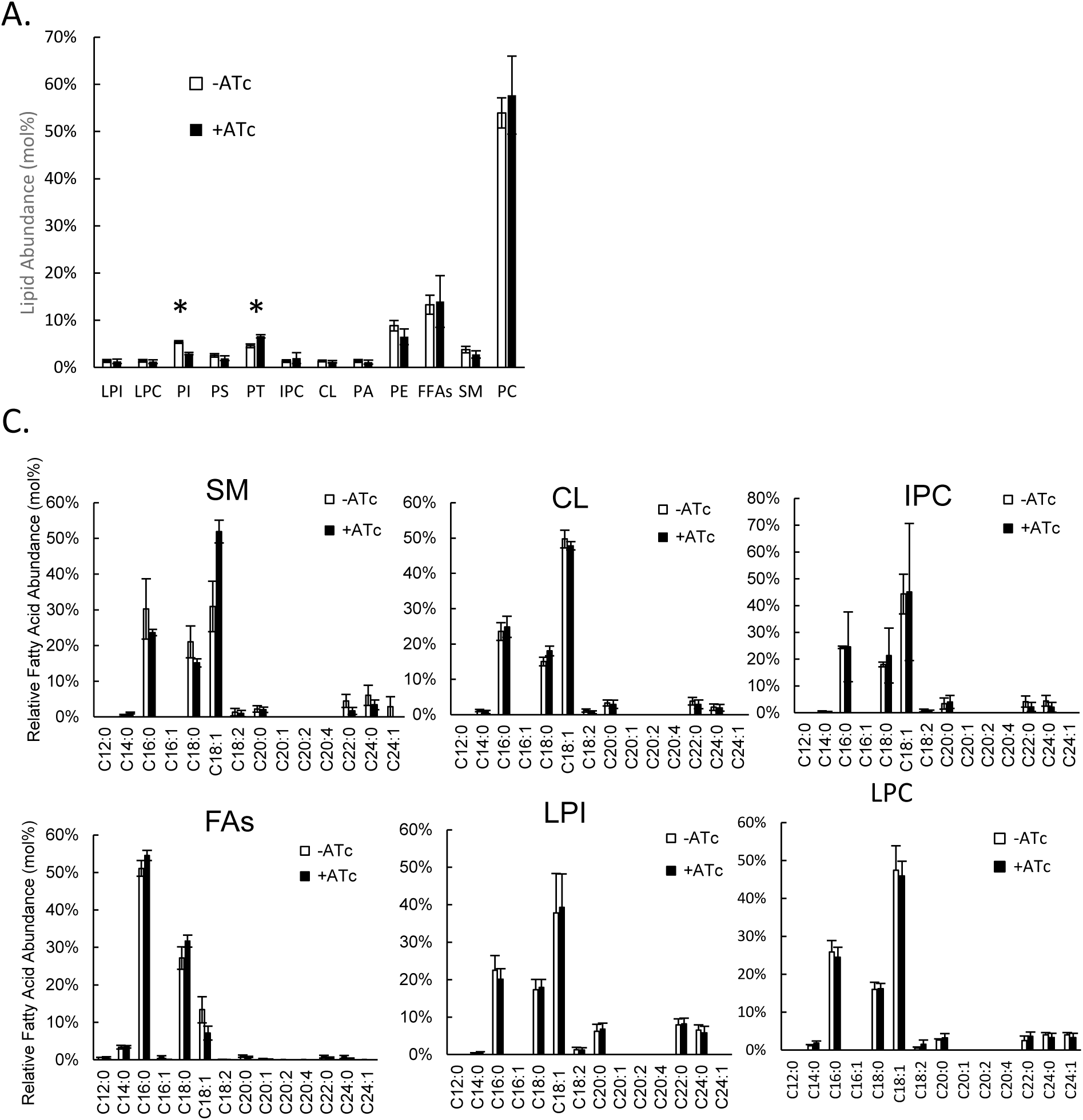
A. Global phospholipid analysis of all phospholipid species.Total lipids expressed as mol%, normalized to total fatty acids. B. Fatty acids profile of Sphingomyelin (SM), Cardiolipin (CL), Inositol Phosphoceramide (IPC) Free Fatty acids (FAs), Lysophosphoinositol (LPI) and Lysophosphocholine (LPC). For all graphs, n=3 biological replicates, error bars represent SEM.

**Figure S4:**
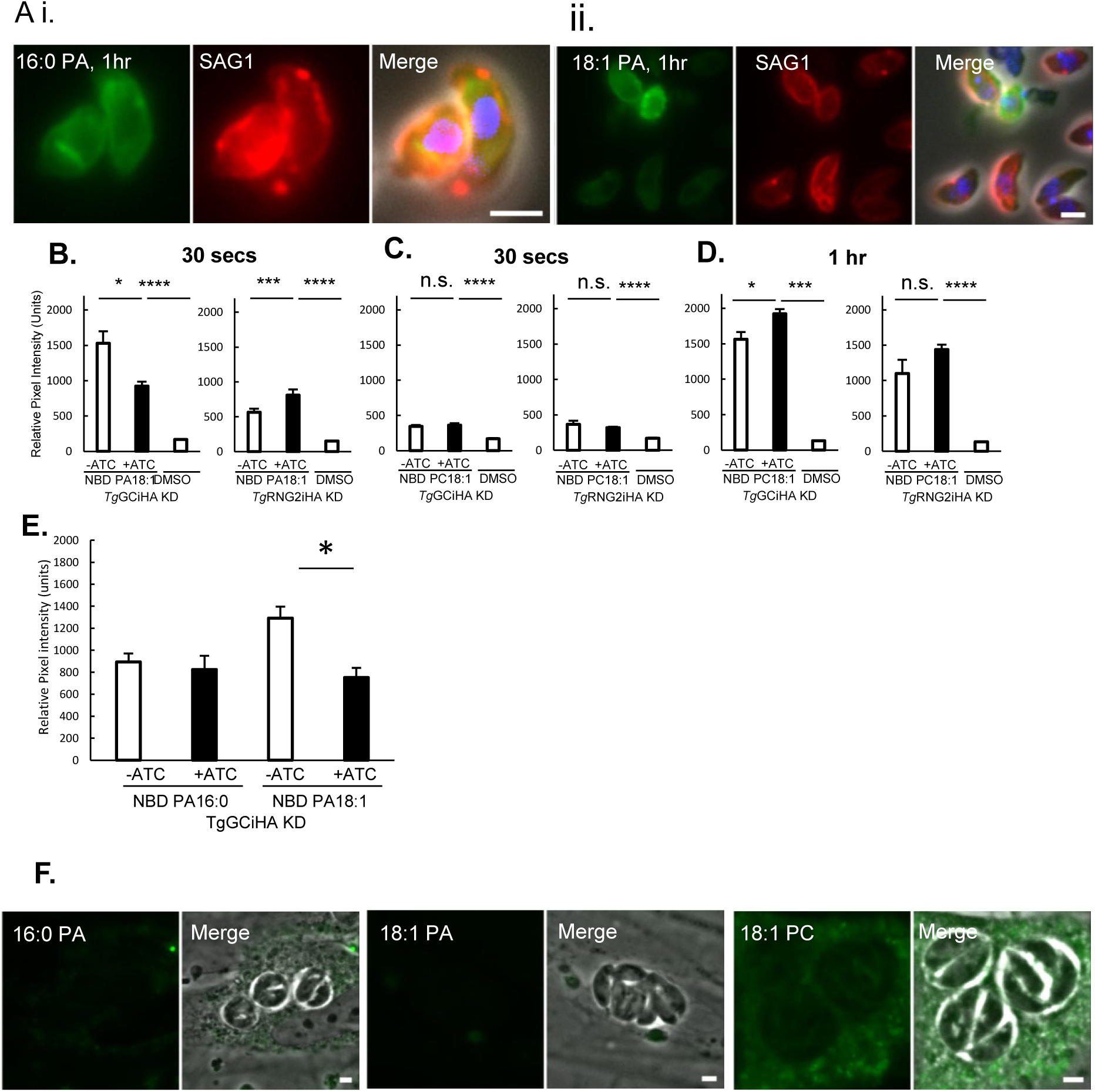
Extracellular *Toxoplasma* tachyzoites readily take up PA. **A.** Lipid uptake assay in extracellular *Toxoplasma* tachyzoites analysed by fluorescence microscopy. Extracellular parasites were incubated in DAG buffer with 5µM of indicated fluorescent NBD-lipid over 1 hour.. **B**. Quantification of fluorescent NBD-PA18:1 uptake by microscopy in TgGCiKD and TgRNG2iKD mutants pre-treated with or without ATc over 30 seconds. n=12 biological replicates **C**. Quantification of fluorescent NBD-PC18:1 uptake by microscopy in TgGCiKD and TgRNG2iKD mutants pre-treated with or without ATc over 30 seconds. n=6 biological replicates **D**. Quantification of fluorescent NBD-PC18:1 uptake by microscopy in TgGCiKD and TgRNG2iKD mutants pre-treated with or without ATc over 1 hour. n=6 biological replicates **E**. Comparison of uptake of fluorescent NBD-PA18:1 and NBD-PA 16:0 by microscopy. n=3 biological replicates **F**. Lipid uptake of intracellular tachyzoites with 5µM of fluorescent NBD-lipids over 24 hoursFor all graphs, \**x* <0.05, ** *x* <0.01, *** *x* <0.005, **** *x* <0.001. All error bars represent SEM. Scale bars represent 2 µm

**Figure S5:**
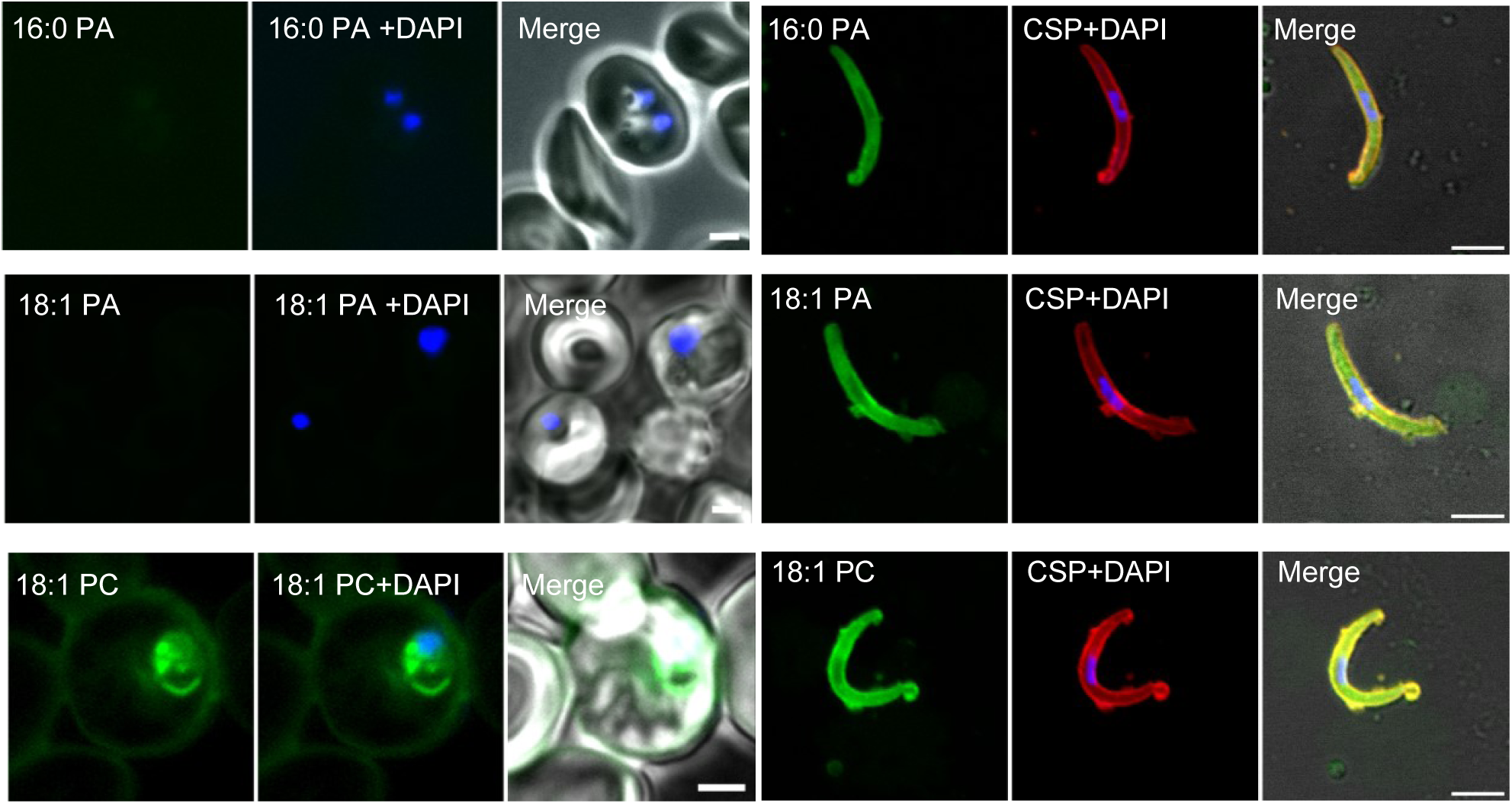
Phospholipid uptake in *Plasmodium falciparum* parasites. **A.** Fluorescent lipid uptake in blood stage *P. falciparum incubated with 5*µM of NBD-PA16:0, NBD-PA18:1, or NBD-PC18:1 for 24 hours. Scale bars represent 2µm **B**. Fluorescent lipid uptake in extracellular sporozoites incubated with 5µM of NBD-PA16:0, NBD-PA18:1, or NBD-PC18:1 for 10 minutes. Scale bars represent 5µm.

**Figure S6:**
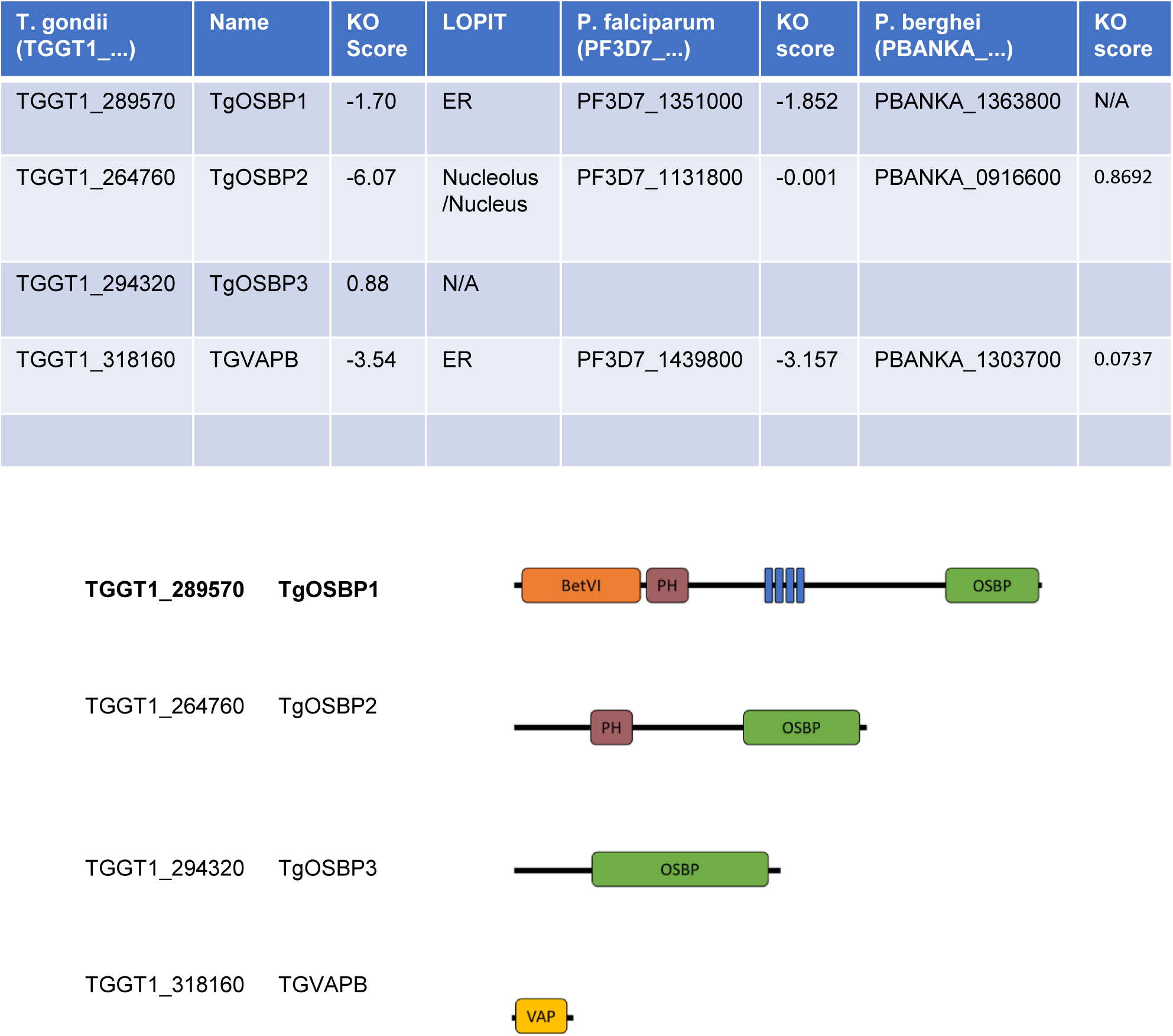
Phylogenetic analysis of genes in the Phosphatidylinositol cycle. **A.** Table of putative OSBPs and VAP cofactors in *Toxoplasma* and *Plasmodium* with predicted hyper-LOPIT localization and essentiality prediction. **B**. Conserved domains of putative OSBP proteins in Toxoplasma and VAP cofafctor.

**Figure S7:**
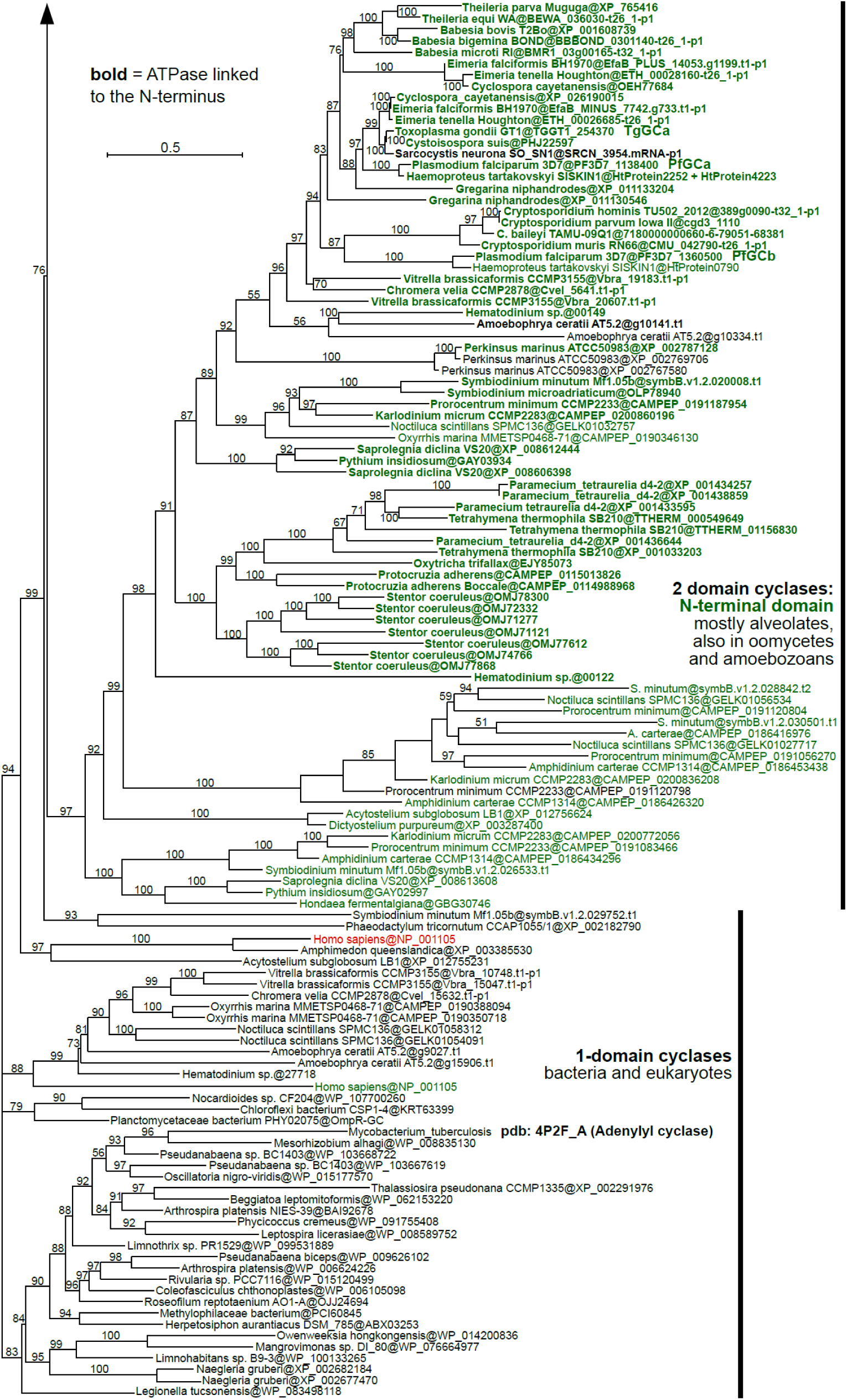
Phylogenetic analysis of N-terminal portion of double cyclase domain of GC. Phylogeny of the cyclase domain (233 protein sequences). Best Maximum Likelihood tree (IQ-TREE) is shown with ultrafast bootstraps at branches. Two-domain cyclase were split in between the N-terminal (in green) and C-terminal somain (in red) and both regions were included in the final alignment. Sequences containing an upstream ATPase domain are shown in bold. Species names are followed by sequence accessions after the @ sign (see Materials and Methods for sequence database sources).

**Figure S8:**
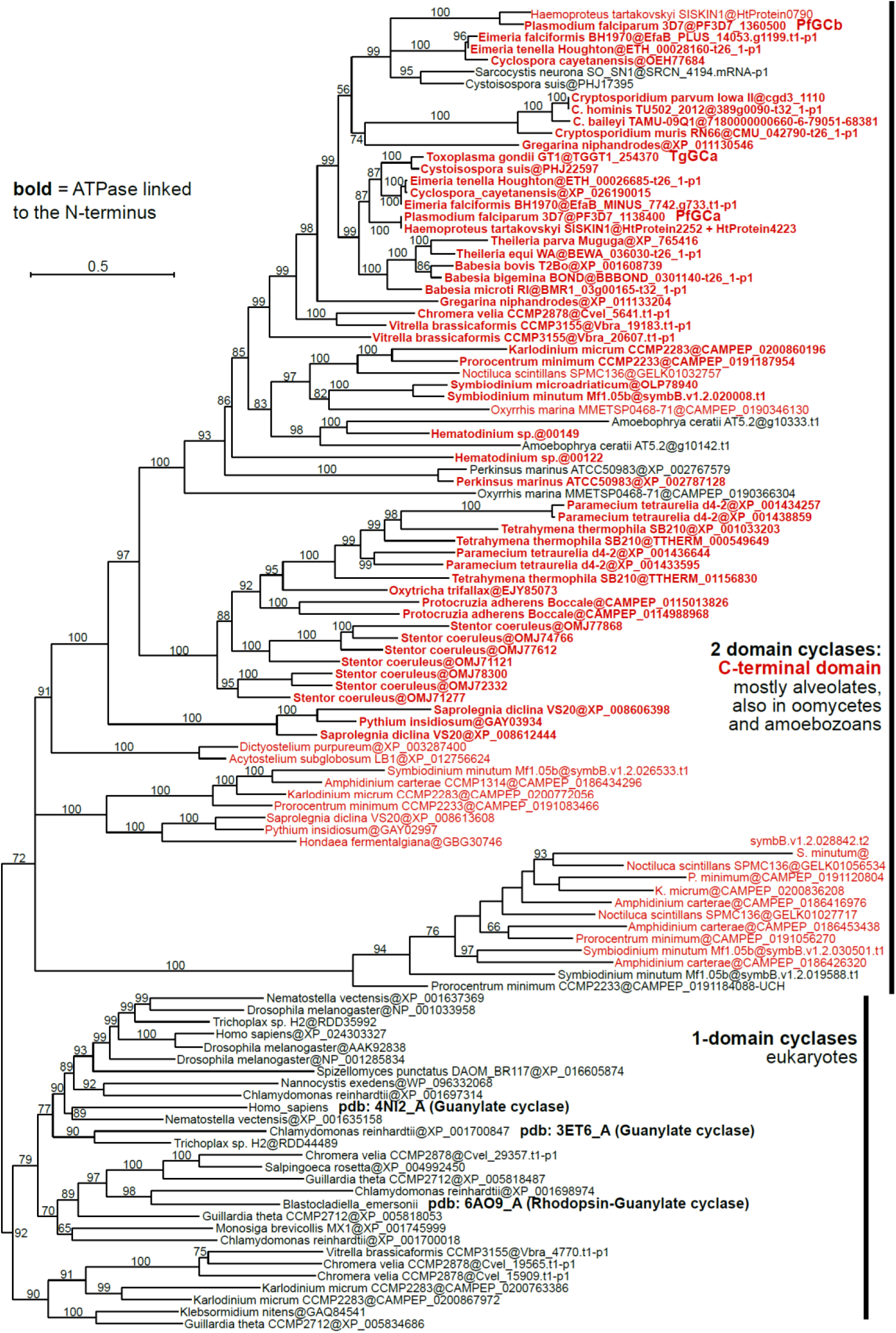
Phylogenetic analysis of C-terminal portion of double cyclase domain of GC. Phylogeny of the cyclase domain (233 protein sequences). Best Maximum Likelihood tree (IQ-TREE) is shown with ultrafast bootstraps at branches. Two-domain cyclase were split in between the N-terminal (in green) and C-terminal somain (in red) and both regions were included in the final alignment. Sequences containing an upstream ATPase domain are shown in bold. Species names are followed by sequence accessions after the @ sign (see Materials and Methods for sequence database sources).

**Figure S9:**
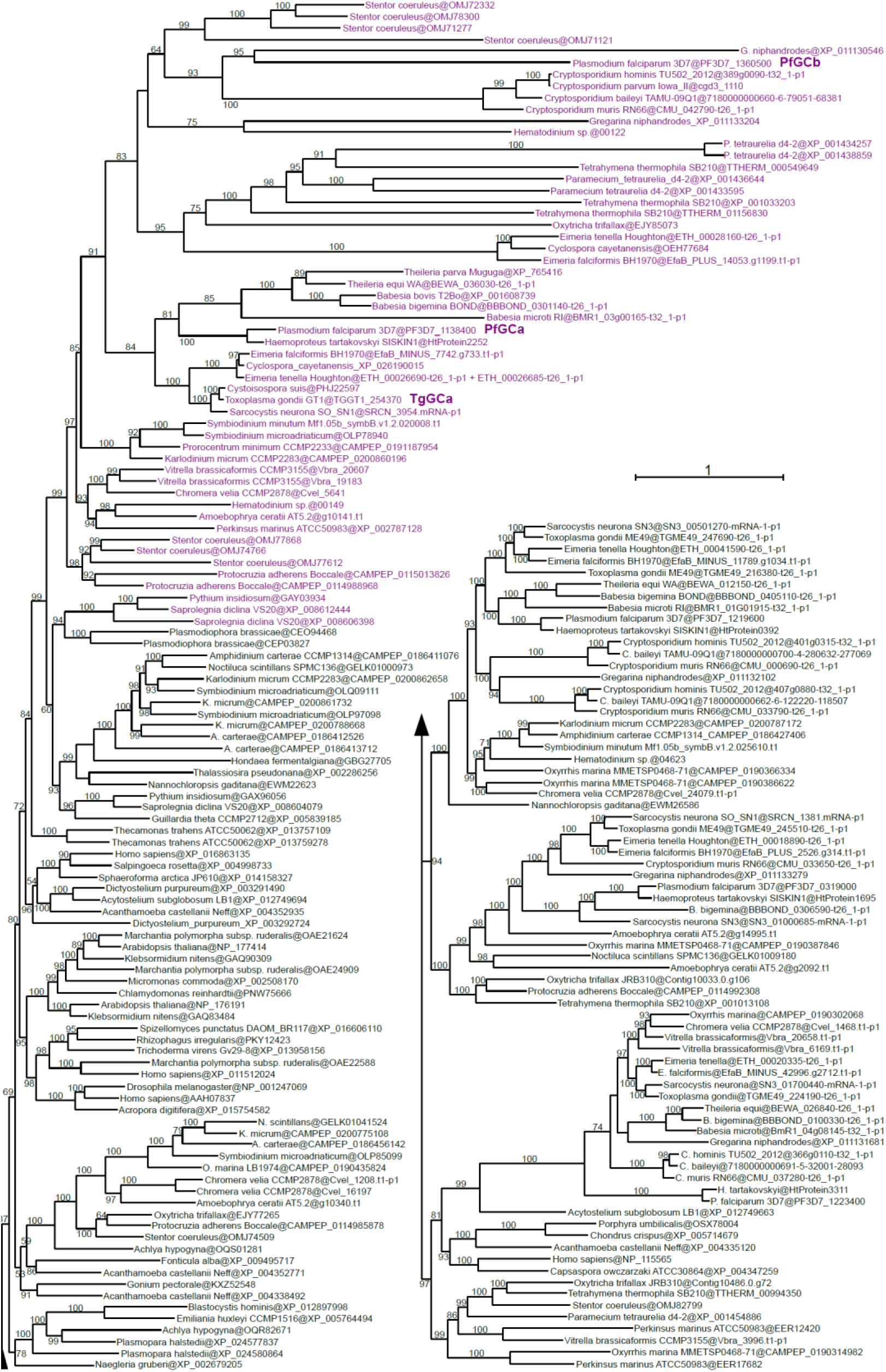
Phylogenetic analysis of putative phospholipid flippases in Alveolates. Phylogeny of DAG kinases (DGKs; 149 protein sequences). Best Maximum Likelihood tree (IQ-TREE) is shown with ultrafast bootstraps at branches. The presence of positively predicted signal peptides is shown by “SP” letters. Species names are followed by sequence accessions after the @ sign (see Materials and Methods for sequence database sources).

**Figure S10:**
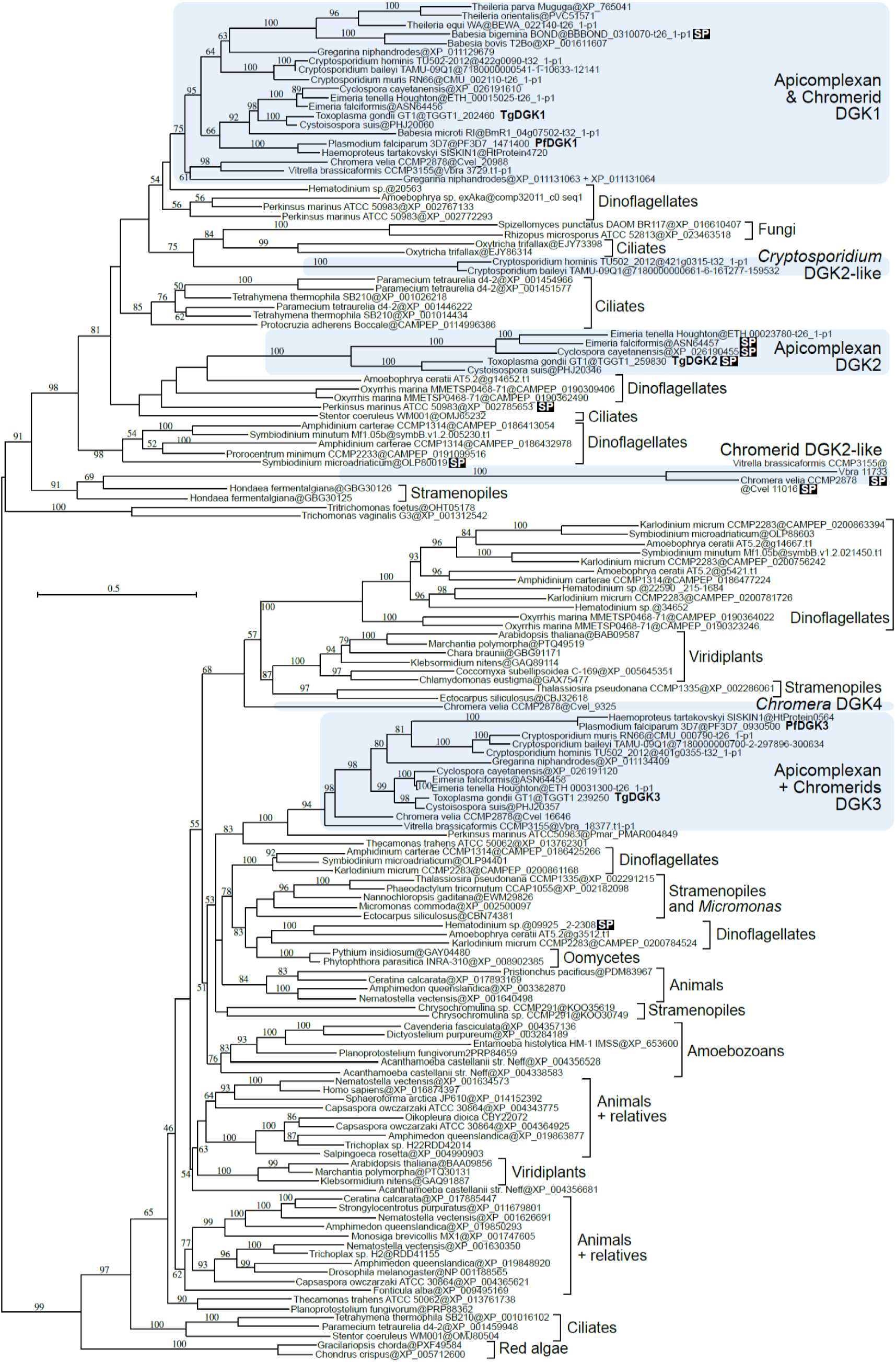
Phylogenetic analysis of DAG Kinases in Alveolates.

**Figure S11:**
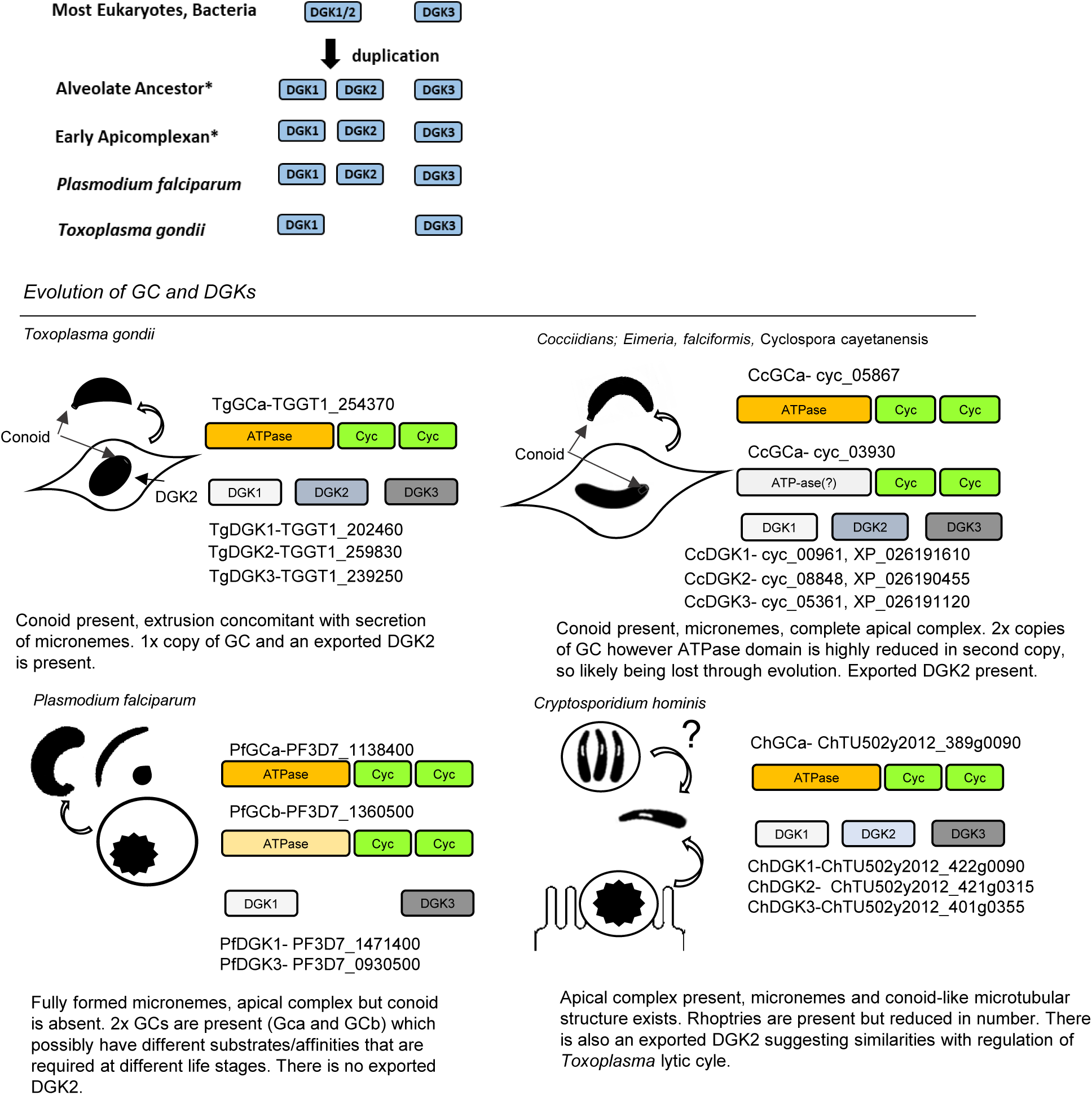
**A.** Evolution of DGK in Toxoplasma and Plasmodium parasites.
**B**. Evolution of GCs and DGKs in Apicomplexan parasites. Coccidia appear to be losing the second copy of GC present in other Apicomplexa. Plasmodium is only species listed here which has lost exported DGK2,

